# Single-Molecule DNA Footprinting and Transcription Imaging Reveal the Molecular Mechanisms of Promoter Dynamics

**DOI:** 10.1101/2025.11.26.690466

**Authors:** Vera Slaninova, Flavia Mazzarda, Lasha Dalakishvili, David Depierre, Christina J. I. Moene, Rachel Topno, Jean-Marc Escudier, Marie-Cécile Robert, Olivier Cuvier, Arnaud Krebs, Nacho Molina, Ovidiu Radulescu, Edouard Bertrand

## Abstract

Live cell RNA imaging revealed that transcription levels are encoded by the intrinsic dynamics of promoters. However, capturing both kinetic and molecular aspects of promoter fluctuations has been challenging. Here, we resolve this key issue by combining Single Molecule DNA footprinting (SMF) with live transcription imaging. Using HIV-1 as a model, SMF reveals that the promoter functions in two modes depending on the viral transactivator Tat. Without Tat, a nucleosome occupies the core promoter and prevents assembly of the pre-initiation complex. With Tat, this nucleosome is absent while TBP and initiating polymerases are frequently detected. Combining live imaging with SMF provides a mechanistic model of promoter dynamics, which estimates the rates of deposition and removal of promoter nucleosomes (0.7 h^-1^), TBP binding (0.04 min^-1^) and polymerase loading (seconds). The data further reveal a kinetic proofreading mechanism of initiating polymerases, which enables Tat to indirectly control promoter nucleosomes by promoting elongation.

## INTRODUCTION

Live cell imaging of transcription using the MS2 or other RNA labeling systems provides a real-time readout of the activity of single promoters ^1^. This showed that transcription is a discontinuous process at steady state, occurring in random bursts of activity that produce several RNAs in a short time, separated by periods of inactivity ^2^. These fluctuations are governed by the intrinsic dynamic properties of promoters, which constantly and stochastically switch between active and inactive states (reviewed in ^1,3–5)^. The switching rates determine the overall promoter strength and the total transcriptional output. Characterizing the molecular and kinetic aspects of promoter dynamics is therefore essential to understand how transcription is controlled.

Transcription imaging has revealed that promoter activity fluctuates on multiple time scales, indicating the existence of many promoter states ^6,7^. Electron microscopy imaging of single promoters and single molecule DNA footprinting (SMF) showed that indeed, promoters occur in different molecular states in different cells of the same population ^8–10^. This diversity confirmed that transcription initiation is a dynamic process in which a promoter constantly switches between different molecular states, some being competent for transcription and others not. This dynamic is driven by the inherent stochasticity of molecular processes operating on a single molecule of promoter DNA. Perturbations experiments have revealed that various events can contribute to the stochastic promoter switches, including enhancer-promoter contacts, chromatin modifications, nucleosomes and transcription factors (TFs) binding and dissociation, pre-initiation complexes stability, promoter-proximal pausing and DNA supercoiling (reviewed in ^1,3–5,11,12^). However, the vast diversity of processes that can potentially influence promoter dynamics, and the wide range of promoter states observed at the molecular level, far exceeds the limited number of states that can be kinetically defined using transcription imaging (typically 2-4 states; ^1,3–5,11,12^). Consequently, the kinetic states are poorly defined at the molecular level. They are however the rate limiting ones, and their molecular characterization is thus essential for a mechanistic understanding of transcriptional control. More generally, while a vast body of the literature indicates that several initiation steps are key points of control ^13–18^, including the deposition/displacement of promoter nucleosomes, the assembly of the pre-initiation complex (PIC) and the release of paused polymerases, the rates of these steps are not well characterized. Addressing this key issue requires methods able to simultaneously integrate molecular and kinetic aspects of promoter dynamics, a challenge that we address here by combining live transcription imaging with SMF.

In addition to determining overall promoter activity, transcriptional fluctuations also contribute to the cell-to-cell variability of gene expression. These can have dramatic phenotypic consequences ^19,20,21^, as highlighted by the case of HIV-1 ^22–24^ (see ^25^ for a review). Antiretroviral therapies can control HIV infections but fail to clear the virus from patients. This persistence is due to latently infected cells, which remain virally silent but can exit latency and produce viral rebounds (reviewed in ^26^). HIV-1 latency is primarily transcriptional and is regulated by a positive feedback loop driven by the viral protein Tat ^26,27^. In the absence of Tat, viral transcription remains low, keeping the virus latent, whereas Tat expression triggers the positive feedback loop, amplifying viral transcription by 100-1000 fold. Latency exit can be triggered by the external stimulation of latent T cells or the spontaneous, stochastic activation of the viral promoter ^25^, both of which then activate the Tat feedback loop ^25–27^. In the case of external stimuli, it has also been shown that latent viruses respond stochastically, with large cell-to-cell heterogeneity even in clonal populations or cells derived from patients ^22,28^. This greatly limits the effectiveness of therapeutic strategies aiming at activating or repressing the latent viral reservoir ^26,25^. Understanding the origin of transcriptional fluctuations is therefore fundamental for HIV-1 biology and therapeutic developments.

Part of the intrinsic dynamic of promoters is genetically encoded in their sequences. The HIV-1 promoter contains a TATA box and binding sites for different transcription factors, including SP1 and NF-kB ^25^. SP1 is necessary for basal and activated transcription, whereas NF-kB mediates the viral response to extracellular signals ^29^. In the absence of Tat, viral transcription is inefficient because of promoter-proximal pausing ^30^, a process in which initiating polymerases transcribe the first 60 to 80 nucleotides and then stop. Pausing is induced by NELF (Negative elongation factor) and DSIF (DRB-sensitivity inducing factor) ^31,32,33^. It is also favored by a nucleosome positioned just downstream of the pausing site (Nuc-1), which acts as a road block ^25–27^. Tat releases the pause by binding to the nascent viral TAR RNA and recruiting P-TEFb, composed of Cyclin T1 and the kinase CDK9 ^25–27^. Tat and Cyclin T1 bind to the TAR RNA and to each other, leading to the formation of a high-affinity complex on nascent viral RNAs ^34^. P-TEFb can also be recruited as part of the SEC complex, whose AFF1/AFF4 subunits improve P-TEFb association with TAR and its kinase activity ^35,36,37^. Once recruited, P-TEFb phosphorylates NELF and DSIF, triggering the dissociation of NELF from the polymerase and the conversion of DSIF into a positive elongation factor ^25–27^. P-TEFb also phosphorylates RNA polymerase II (RNAPII) on the Serine 2 of its C-terminal domain (CTD) and on the linker between the polymerase core and the CTD ^33,38^. This promotes the binding of SPT6 and the stimulation of transcription elongation ^33,38^. Finally, in the case that a paused polymerase fails to enter elongation, it is removed from DNA by the Integrator complex, which cleaves the nascent RNA and induces termination ^39,40^.

Interestingly, Tat also plays a role in remodeling the chromatin of the HIV-1 promoter ^41–43^. In the absence of Tat, the promoter harbors two well positioned nucleosomes termed Nuc-0 and Nuc-1 ^25,43^. Nuc-0 is located approximately 250 nt upstream of the TSS whereas Nuc-1 lies just downstream the TSS and contributes to transcriptional repression ^44^. These nucleosomes are separated by a DNase hypersensitive region (DHS1) thought to be devoid of nucleosomes, leaving accessible the core promoter with the TATA box, the NFkB and SP1 binding sites ^25,43^ In presence of Tat, promoter nucleosomes become more acetylated and Nuc-1 is repositioned to increase the accessibility of the transcription start site (TSS) ^45,46^. This involves the histone acetyl transferases p300/CBP and p/CAF, and the nucleosome remodeling complexes BAF and P-BAF ^41,42,47^, but how Tat affects the rates of nucleosome deposition and removal is still not well understood despite their importance in viral transcription.

HIV-1 transcription has been previously imaged in live single cells using the MS2/MCP-GFP system ^6,48^. This revealed that viral transcription fluctuates stochastically in the presence and absence of Tat, and that the promoter responds to latency reversing agents with large cell-to-cell differences. This data enabled the development of kinetic models describing active and inactive promoter states, but have fallen short of defining their molecular nature ^6,48^. For instance, while binding and dissociation of nucleosomes and TBP, as well as polymerase pausing, are known to be key steps regulating viral transcription ^25–27,43,45^, their rates, and how they are modulated in different conditions, remains largely unknown. Here, we integrated live transcription imaging with single molecule DNA footprinting to resolve this fundamental issue. Indeed, the two methods offer complementary insights into transcription mechanisms: SMF reveals the number, occupancy probabilities, and molecular nature of discrete transcriptional states, while MS2 imaging provides dynamic information on the stochastic transitions between them. By modeling together data from SMF and live transcription imaging, we provide, for the first time, a molecular description of the dynamics of a promoter. This reveals the rates of key steps of transcription initiation and how they are regulated. It also uncovers an important kinetic proofreading mechanism controlling transcription initiation.

## RESULTS

### SMF reveals the molecular occupancy of the HIV-1 promoter

We aimed to use single molecule DNA methylation footprinting (SMF; ^10,49^) to simultaneously quantify nucleosomes, RNAPII and TFs occupancy on single molecules of the HIV-1 promoter (Fig. 1A), thereby characterizing the diversity of its molecular states. We previously developed cell lines bearing an HIV-1 reporter optimized for live cell imaging of transcription with the MS2/MCP system (Fig. 1B; ^50,51^). The reporter contains an intronic array of 128 MS2 stem-loops downstream the HIV-1 LTR, which contains the viral promoter (Fig. 1C). It was inserted at a specific locus in HeLa cells and isogenic cell lines expressing a saturating amount of Tat (‘High Tat’ cells), or lacking Tat (‘No Tat’ cells) were created, mimicking active and latent viral states ^6,48^. Consequently, these cells express the HIV-1 reporter at widely different levels ^48^. We took advantage of these existing cell lines to perform the SMF analysis. Nuclei were isolated and incubated with the M.CviPI enzyme that methylates all accessible GpCs. Binding of RNAPII, TFs or nucleosomes prevents methylation and therefore leaves a characteristic footprint in each DNA molecule ^10,49^. Following DNA extraction and bisulfite conversion to transform unmethylated Cs into Ts while methylated Cs remain Cs, the sequences of the HIV-1 promoter and control genomic regions were PCR amplified and deep-sequenced (note that the HIV-1 amplicon covers the -336/+164 region with respect to the transcription start site; see Fig. 1C). This enables the determination of the methylation status and protection patterns for all the relevant cytosines, for each molecule of the HIV-1 promoter present in the original population (Fig. 1A).

**Figure 1:**
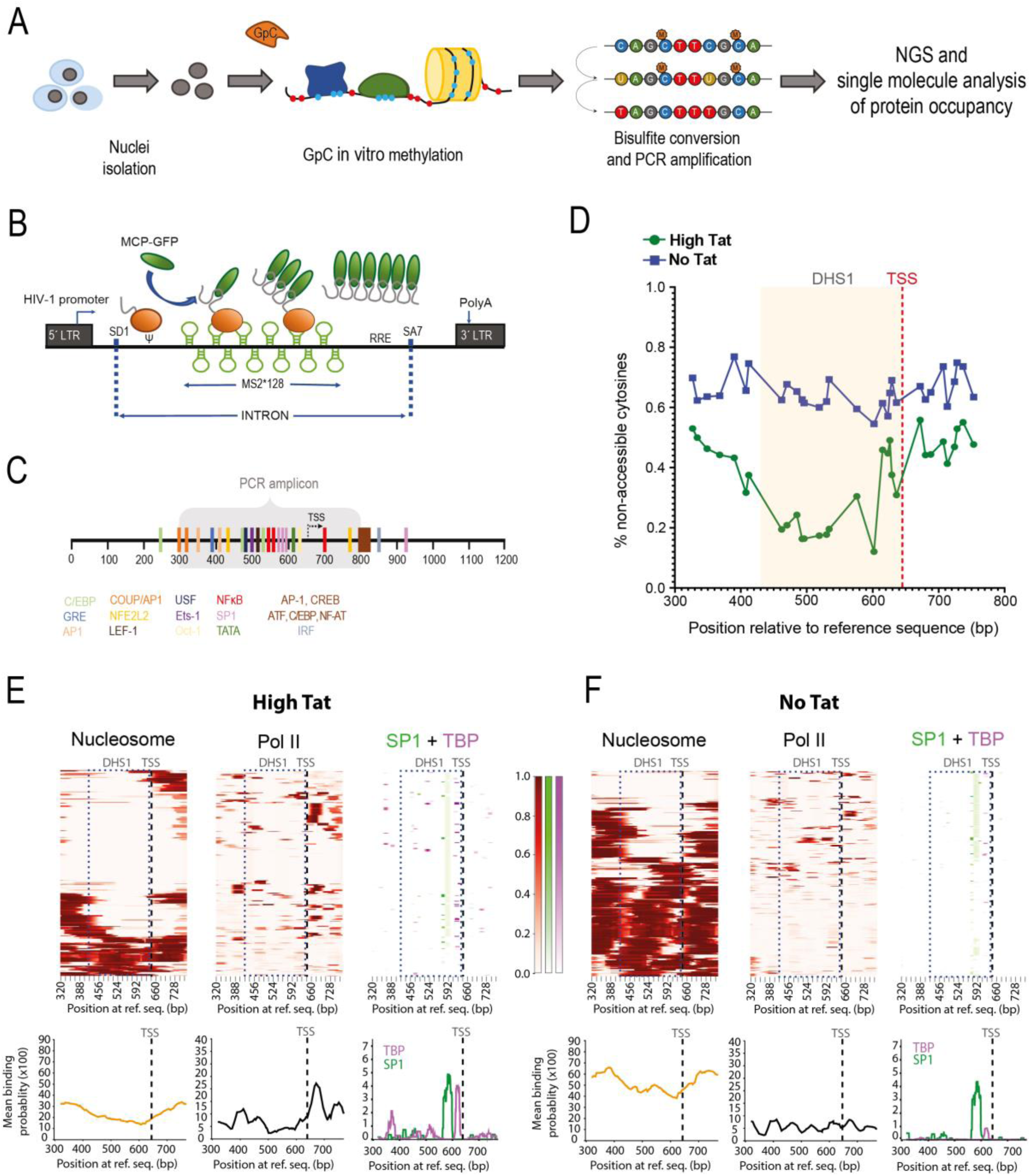
SMF reveals the molecular diversity the HIV-1 promoter. **(A)** Scheme of SMF pipeline: isolated nuclei are treated *in vitro* with a GpC methyltransferase. Extracted DNA is bisulfite converted and regions of interest are enriched using PCR amplification. Resulting libraries are subjected to Illumina sequencing. **(B)** Schematic of the HIV-1 reporter. SD1: major HIV-1 splice acceptor, SA7: last HIV-1 splice site acceptor, Ψ: packaging signal, RRE: Rev-responsive element, LTR: long terminal repeat. Green loops represent MS2 repeats, the orange ball is RNAP II (with nascent mRNA), and green ovals are MCP-GFP. **(C)** Schematic of the HIV-1 5’LTR with known TF binding sites (color-coded). The area amplified for SMF is highlighted in grey. **(D)** Average accessibility of the GpC dinucleotides of the HIV-1 promoter in ‘High Tat’ (green) and ‘No Tat’ (blue) cells. The yellow area marks the DHS1 region and the red dashed line marks the TSS. Plotted values are the mean of four (No Tat) and three (High Tat) independent experiments. **(E-F)** Binding probabilities of nucleosomes, RNAP II and various transcription factors in High Tat **(E)** and No Tat **(F)** cells. Probabilities are estimated by Hiddenfoot. Top panels: each lane is a single molecule, the blue rectangle indicates the DHS1 area, the black dashed line marks the TSS and the color scale bars give the binding probability. Bottom panels: average binding probabilities for the indicated factors. SP1 is in green and TBP in purple.

An initial analysis of the bulk accessibility profiles, representing the averages of many molecules, showed that the majority of the HIV-1 promoters were globally protected in absence of Tat (each target cytosine was accessible in only 20 to 40% of the promoters), indicating a poorly accessible chromatin likely covered by nucleosomes. In contrast, the promoter was much more methylated in presence of Tat (40-85% of accessibility for each target cytosine), indicating a more open chromatin (Fig. 1D). To control for experimental variability, we used two genomic regions encompassing CTCF binding sites located near the promoters of TP53 and ABL2. SMF analysis showed no differences in the accessibility of the control regions between ‘High Tat’ and ‘No Tat’ cells, indicating that the effects of Tat are specific for the HIV-1 promoter (Fig. S1A).

To generate high-resolution mapping of nucleosomes, TFs, and RNAPII on single promoters, we applied HiddenFoot, a biophysical modeling approach that infers probabilistic binding profiles from DNA methylation footprints ^52^. Briefly, HiddenFoot integrates known TF binding motifs and uses their position weight matrices (PWMs) to compute sequence-specific binding affinities. To infer the footprints, the model assumes that bound TFs protect DNA segments from methylation based on the lengths specified by their PWMs. For RNAPII and nucleosomes, the model assumes protection lengths of 40 and 147 nucleotides, respectively, and sequence-independent binding. Using a thermodynamic framework, HiddenFoot then evaluates all possible binding configurations of TFs, RNAPII, and nucleosomes on each sequenced DNA molecule, and estimates the probability of occupancy for each factor at every nucleotide position. To analyze TF binding on the HIV-1 promoter, we ran HiddenFoot with 15 PWMs from available databases ^53,54^, corresponding to TFs with high-quality binding sites within the HIV-1 promoter, which included NF-kB, SP1 and TBP (Fig 1C; see Methods).

We grouped the Hiddenfoot results into three maps describing the probability of binding of nucleosomes, RNAPII and TFs, providing single molecule profiles and averages (Fig. 1E and 1F). A striking difference between ‘High Tat’ and ‘No Tat’ cells was the frequent detection of nucleosomes in the absence of Tat. Nucleosomes were detected over the entire promoter sequence, including the DHS1 region that covers the core promoter. Our data indicate that the average probability of detecting a nucleosome over DHS1 was 0.45-0.55 in ‘No Tat’ cells, which was in stark contrast to ‘High Tat’ cells where it was ∼0.15 (Fig. 1E and 1F, bottom panels). This result was unexpected because previous work indicated a nucleosome-free region over the DHS1 area, even in the absence of Tat (see introduction; ^55^), even if one study observed a labile DHS1 nucleosome on a latent HIV-1 virus ^47^. Importantly, the DHS1 nucleosome, thereafter termed Nuc-DHS1, is positioned over the binding sites of SP1, NF-kB and TBP. It therefore likely plays an important regulatory role by preventing these factors from binding, thereby limiting transcription initiation in absence of Tat.

The HIV-1 promoter in ‘High Tat’ cells had frequent RNAPII footprints in the +1 to +120 region. Similarly, footprints of the key transcription factors SP1 and TBP could be detected in ‘High Tat’ cells. Although SP1 binding remained similar in presence and absence of Tat (Fig. 1E), as previously observed ^56^, the binding probability of TBP dropped ∼6 fold in the ‘No Tat’ cells (Fig. 1E and 1F). This was consistent with the lower transcriptional activity observed in these cells, as well as with previous TBP ChIP data ^56^. Promoter proximal pausing of RNAPII is a key regulatory step controlled by Tat, with frequent pausing occurring in its absence. However, we found that only few promoters had a promoter-proximal RNAPII footprint in absence of Tat (Fig. 1F; Fig. S7), suggesting that pausing alone is probably insufficient to account for the low transcriptional activity of cells lacking Tat. Taken together, SMF revealed two unexpected features of the ‘No Tat’ cells: a high nucleosome occupancy of the DHS1 region covering the core promoter, and a low frequency of promoter-proximal polymerase footprints.

### Validation of the footprints

To ensure that the SMF binding probability maps inferred by HiddenFoot reliably reflected molecular occupancies, we generated ‘High Tat’ cells with point mutations in the HIV-1 promoter, in the TATA box and one of the SP1 binding sites (Fig. 2A and 2B). Measurement of nascent HIV-1 RNAs by smFISH indicated that these mutations led to a decreased viral transcription (∼2 fold for SP1 and ∼8 fold for the TATA box; Fig. S2A). We then performed SMF and compared the results to the wild-type promoter. Bulk methylation footprinting data showed a drop in SP1 and TPB footprints in their respective binding site mutants (Fig. 2C and 2D). Likewise, single molecule analysis showed that the average binding probability of SP1 dropped from ∼0.05 in the wild-type to <0.01 in the SP1 mutant (Fig. 2E and 2G), while the binding probability of TBP decreased from 0.05 in the wild-type to <0.005 in the TATA mutant (compare Fig. 2F and 2H to Fig. 1E). Consistently, the RNAPII footprints diminished in the SP1 and the TATA mutant cells while nucleosome occupancy increased (compare the solid and dashed lines in Fig. 2E, 2F). Again, the internal control genomic regions were similar in all conditions (Fig. S2B).

**Figure 2:**
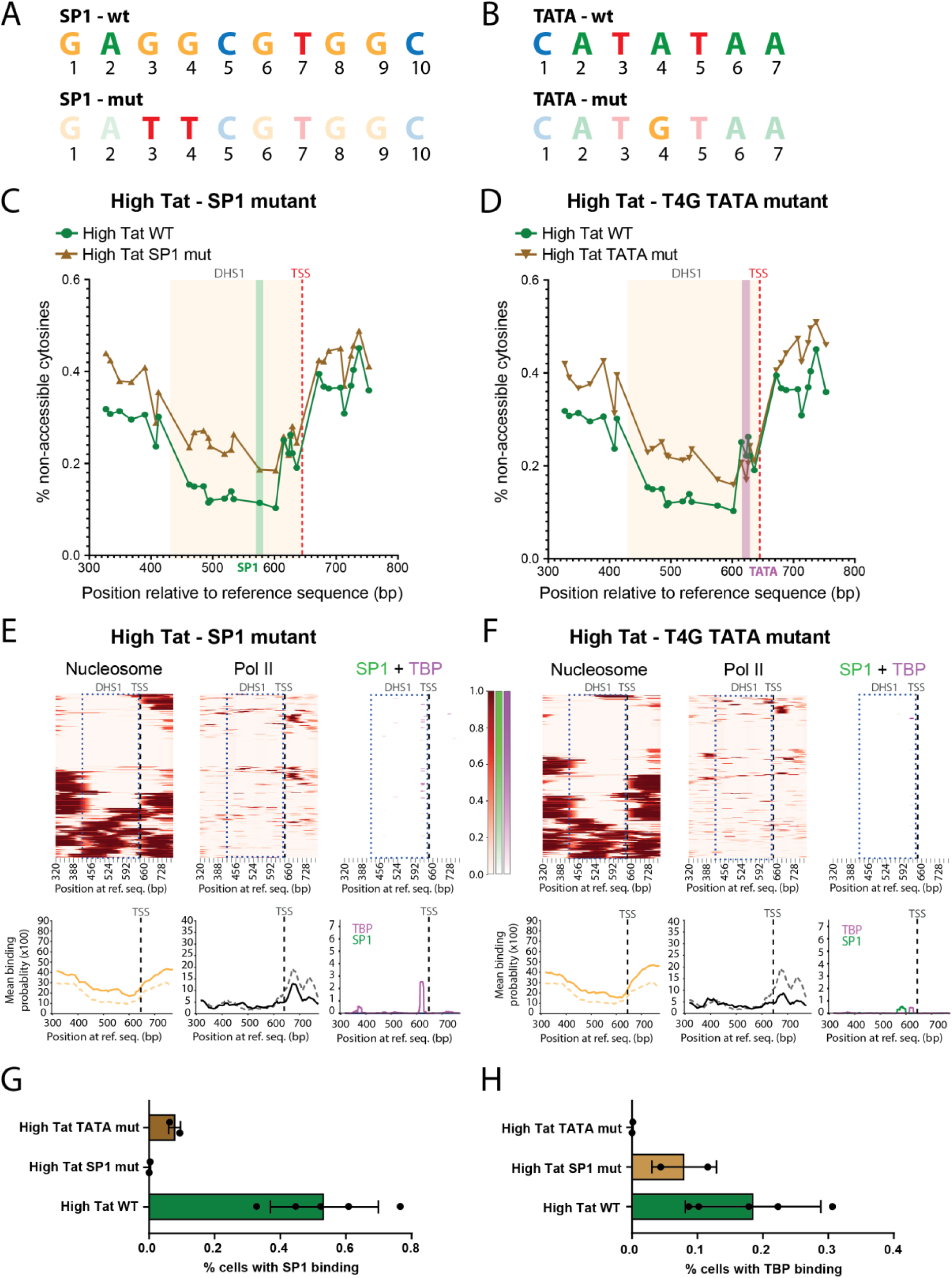
SMF captures the binding of transcription factors to the HIV-1 promoter. **(A-B)** Sequences of HIV-1 SP1 binding site **(A)** and TATA box **(B)** and their respective mutants. **(C-D)** Average accessibility of the GpC dinucleotides of HIV-1 promoter in High Tat SP1 mutant **(C)** and High Tat T4G TATA mutant **(D)** cells, in comparison with High Tat WT cells (green in both panels). The yellow area marks the DHS1 region and the red dashed line marks the TS. Green and purple bars respectively mark the SP1 and TBP binding site. **(E-F)** Binding probability of nucleosomes, RNAP II and various transcription factors in High Tat SP1 mutant **(C)** and High Tat T4G TATA mutant **(D)** cells. Probabilities are estimated by Hiddenfoot. Top panels: each lane is a single molecule, blue rectangle gives the DHS1 area, black dashed line marks TSS, and the color scale bars give the binding probability. Bottom panels: average binding probabilities for the indicated factors. SP1 is in green and TBP in purple. Dashed lines show the mean binding probabilities of nucleosome and RNAP II in High Tat WT cells. **(G-H)** Fraction of promoters occupied at the SP1 binding site **(G)** and the TATA box **(H),** in High Tat SP1 mutant cells, High Tat TATA mutant cells, and the High Tat WT controls. Error bars correspond to the standard deviation and individual data points are indicated. The fraction of occupied molecules is determined by thresholding the single molecule binding probabilities (see text and Fig. S2).

The probability of binding relates to occupancy but the relation between the two is dependent on the abilities of SMF and Hiddenfoot to detect binding. For instance, a factor may be more difficult to detect than another one, yielding low binding probabilities even when bound. To evaluate how occupancy and binding probabilities relate to each other, we plotted the distributions of the binding probabilities of RNAPII, SP1 and TBP, the latter two considering only their cognate binding sites (Fig. S2D-G). Nucleosomes and RNAPII showed a large distribution of binding probabilities, spreading from 0 to 1 for nucleosomes and 0 to 0.5 for RNAPII. In contrast, the distributions of binding probabilities for SP1 and TBP were smaller, spreading mainly from 0 to 0.05, possibly because the shorter binding sites of these factors reduced their ability to be detected. To convert the binding probabilities in molecular occupancies, we overlaid the distribution obtained with the wild-type and mutant reporters. The probability distributions of the SP1 and TBP mutants were skewed toward 0, barely exceeding the first bin. We then binarized the binding probabilities using a threshold that best separated the mutant from the wild-type (Fig. S2D-G). This estimated that 54% and 18% of the promoter molecules were bound by SP1 or TBP in High Tat cells, respectively, as compared to <1% in the corresponding mutants.

Next, we aimed to validate the footprint observed for RNAPII. To this end, we treated ‘High Tat’ and ‘No Tat’ cells with Triptolide, a TFIIH inhibitor that prevents new initiation events (Fig. 3). After treating cells with 1 mM Triptolide for 1h, we observed that in ‘High Tat’ cells, RNAPII footprints after the TSS became virtually undetectable (Fig. 3A and 3B). Interestingly, this was accompanied by a slight increased detection of the Nuc-1 and Nuc-0 nucleosomes that are flanking the core promoter, while the DHS1 region was not affected (Fig. 3A and 3C). As with SP1 and TBP, the single molecule binding probabilities of RNAPII were binarized using a threshold set with the Triptolide treated cells, considered as a negative control. This estimated that ∼40% of the promoters were occupied by RNAPII in untreated ‘High Tat’ cells, and <5% after Triptolide treatment (Fig. 3B, bar graph, Table S1). In ‘No Tat’ cells, the levels of RNAPII footprints located after the TSS diminished after 1h of Triptolide treatment (Fig. 3E). Taken together, these data indicated that the HIV-1 promoter is suitable for SMF analysis and that the probability binding maps faithfully captured the association of RNAPII and the occupancy of the TATA box and SP1 binding sites.

**Figure 3:**
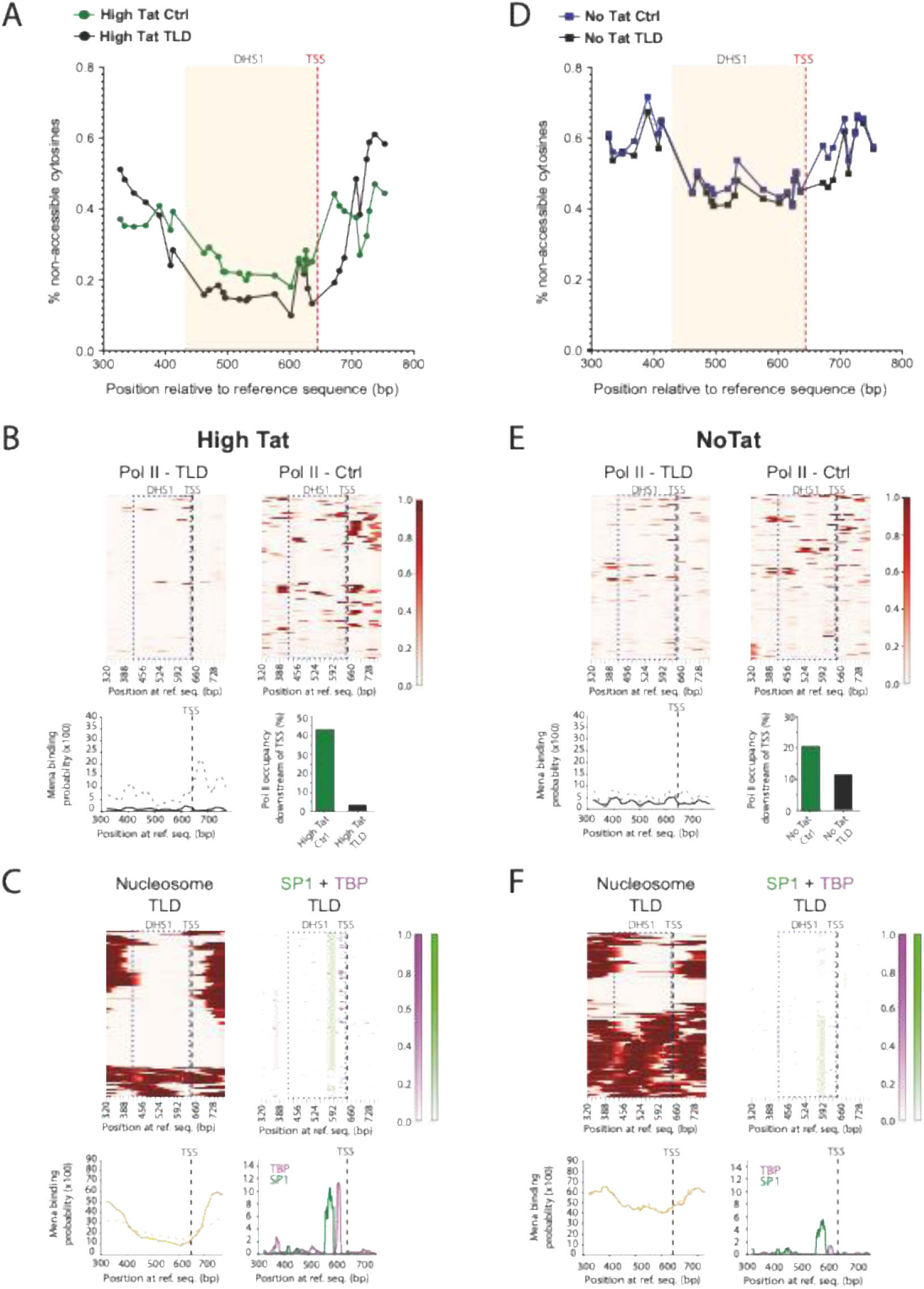
SMF captures the occupation of the HIV-1 promoter by RNAPII. **(A) and (D)** Average accessibility of the GpC dinucleotides of the HIV-1 promoter in High Tat **(A)** or No Tat cells **(D)**, treated with triptolide (black) and compared to non-treated control (green and blue, respectively). The yellow area marks the DHS1 region and the red dashed line marks the TSS. **(B, C) and (E, F)** Binding probabilities of nucleosomes, RNAP II and various transcription factors in High Tat **(B, C)** and No Tat **(E, F)** cells treated with triptolide. Probabilities are estimated by Hiddenfoot. Top panels: each lane is a single molecule, blue rectangle gives the DHS1 area, the black dashed line marks TSS, and the color scale bars give the binding probability. Bottom panels: average binding probabilities for the indicated factors. SP1 is in green and TBP in purple. Dashed lines show the mean binding probabilities of nucleosome and RNAP II in untreated High Tat WT cells. The bar graphs in (B) and E) show the fraction of molecules occupied by RNAPII downstream of TSS, with and without treatment. Occupancy was measured after thresholding the single molecule binding probabilities (see text and Fig. S2).

### Paused polymerases are infrequently detected in absence of Tat

Using SMF, we observed a low detection of RNAPII footprints on HIV-1 promoters in absence of Tat, with an estimate of ∼15% of the promoters being occupied by a polymerase (Fig. 1F, Table S1). This was somewhat surprising because promoter proximal pausing is expected to be frequent in this situation ^25–27^. Because smFISH can reliably detect and count single RNAs, we turned to this method to obtain a second estimate of the fraction of promoters with a paused polymerase. To this end, we developed a triple-color reporter system to reliably detect the nascent TAR RNAs that are characteristic of paused polymerases. Since TAR is only 60 nucleotides long, we sought to improve RNA detection by inserting four MS2 stem-loops directly within the TAR element. To control for RNA folding, we generated two variants, referred to as TAR-MS2*4c2 and TAR-MS2*4c3 (Fig. 4A). In the first design, the TAR element completes its folding after transcribing the MS2 sites, while in the second design, TAR folds independently of the MS2 stem-loops. The two reporters also contained an intronic array of 64 SacB repeats positioned in place of the original intronic MS2*128 sites. Using MS2 and SacB smFISH probes, it was thus possible to visualize both short nascent transcripts attached to the paused polymerase (TAR-MS2 positive, SacB negative), and the elongated RNAs that are produced once the pause is lifted (positive for both TAR-MS2 and SacB). To improve detection, we included 96 TetO repeats ∼1,6 kb upstream of the viral promoter. This way, the HIV-1 insertion site was easily visualized using a TetR-tagBFP expression plasmid, enabling us to precisely point the HIV-1 transcription site within nuclei and unambiguously identify nascent TAR RNAs (Fig. 4B).

**Figure 4:**
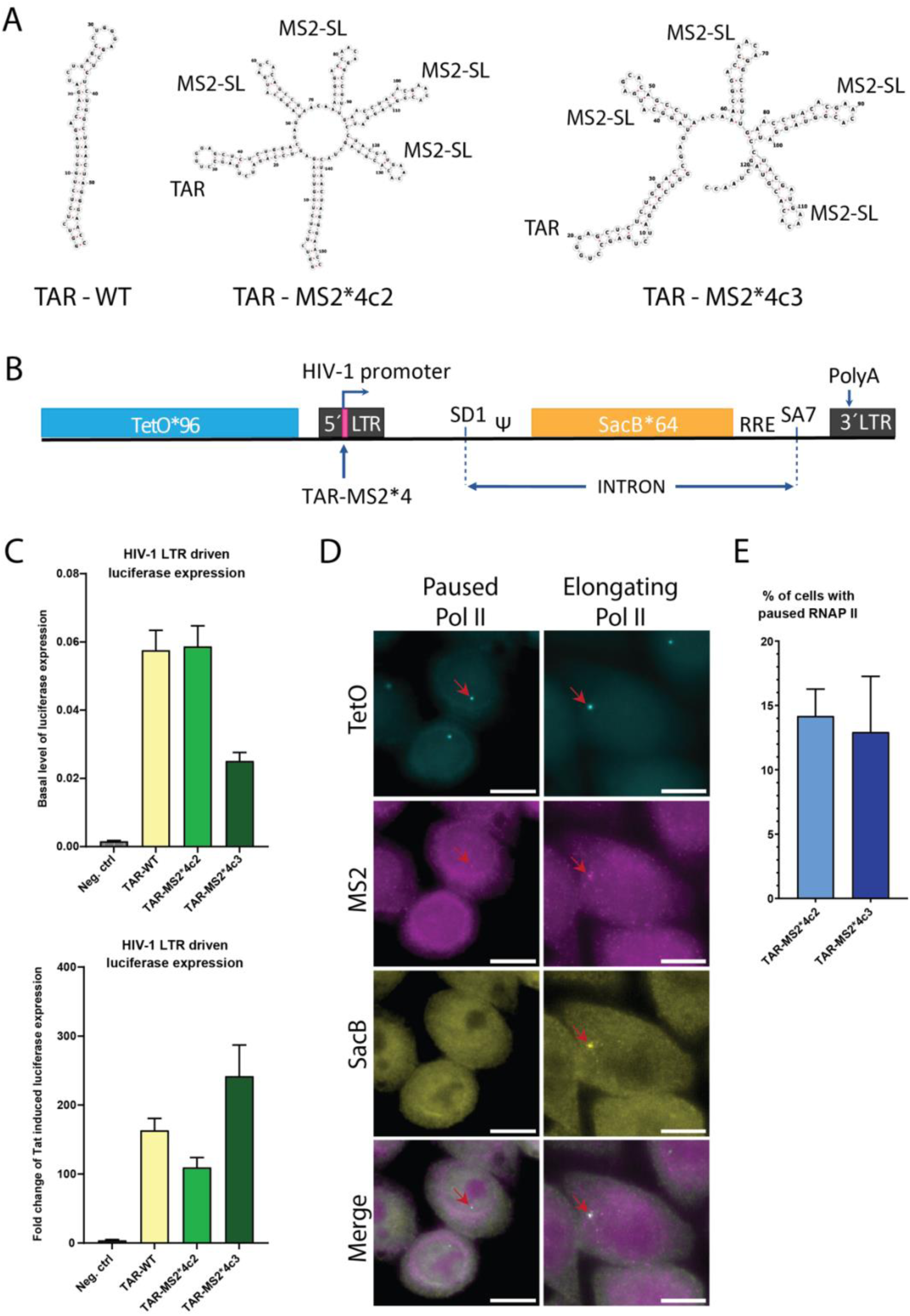
Polymerase pausing is infrequently detected. **(A)** Structure of wild type TAR element (left), TAR-MS2*4c2 (center) and TAR-MS2*4c3 (right). **(B)** Schematic of the HIV-1 triple color reporter system. The 96 TetO repeats (blue) are located upstream of the 5’ LTR to indicate the localization of the reporter gene in the nucleus. Four MS2 stem-loops are inserted into the TAR element (red bar) to allow visualization of the short nascent RNA that are transcribed before pausing occurs. 64 SacB repeats (yellow) are located downstream of the pausing site to visualize the elongated transcripts. **(C)** The bar plots display the luciferase activity of the HIV-1 TAR-MS2*4c2 and TAR-MS2*4c3 LTRs, as compared to HIV-1 WT LTR control, under basal (top) and Tat-induced (bottom) conditions. The values were obtained from extracts of HeLa cells transiently transfected with plasmids having the firefly luciferase under the controls of the LTRs. pBlueScript was used as a negative control (Ctrl). Firefly activities were normalized to that of a co-transfected renilla expression vector to correct for transfection efficiencies. The error bars represent the standard deviations (n>=3). **(D)** Images are micrographs of HeLa cells stably expressing the HIV-1 TAR-MS2*4c2 reported, and labelled for the TetO repeats (cyan, stained with TetR-TagBFP); short nascent HIV-1 RNAs (purple, stained by smFISH with Cy3 TAR-MS2 probes) and long nascent HIV-1 RNAs (yellow, stained by smFISH with Cy5 SacB probes). Scale bar: 10 microns. Arrows point to the TetO array, which indicates the HIV-1 transcription site. **(E)** Quantification of paused RNAPII using the triple color reporter system. Paused polymerases are MS2 positive and SacB negative at the TetO array.

We first verified that the modified TAR elements were functional. For this, the promoters were cloned upstream of firefly luciferase and both the basal and Tat-induced activity of these constructs were measured by luciferase assays (Fig. 4C). Overall, both the TAR-MS2*4c2 and TAR-MS2*4c3 promoters were strongly responding to Tat (100-250 fold activation), similar to the wild-type despite the addition of four MS2 stem-loops into TAR. For imaging, we integrated the constructs into HeLa cells at the same locus that was used for the previous SMF experiments. We then developed sets of 7 and 5 fluorescent smFISH probes ^57^, covering the TAR-MS2*4c2 and TAR-MS2*4c3 sequences, respectively (Fig. S4A), and a set of 39 probes targeting the SacB region (Table S2). The detection of short RNA molecules is inherently difficult due to the small number of probes and the resulting low signal-to-noise ratio. To verify that we were indeed able to reliably detect single molecules of the short TAR transcripts, we counted the nucleoplasmic single RNAs detected by the SacB and TAR-MS2 probes. We observed that the majority (77%) of the single RNAs detected by SacB probes were also labeled by the TAR probes, confirming the efficiency of these probes to detect short nascent transcripts (Fig. S4B).

We then analyzed the smFISH data obtained in the TAR-MS2*4c2 and TAR-MS2*4c3 cell line in the absence of Tat. We counted the number of MS2 and SacB spots colocalizing with the TetO repeats. When the signal was observed only in the MS2 channel, the promoter was classified as “paused”. If both the MS2 and SacB signals were present, the reporter was considered to be in productive elongation (Fig. 4D). The results showed that in both the TAR-MS2*4c2 and TAR-MS2*4c3 cells, ∼15% of the promoters were in the “paused” state, i.e. with TAR-MS2 but no SacB RNAs (Fig. 4E). This was consistent with the SMF analysis, which estimated a similar proportion of promoter-proximal polymerases, confirming that paused polymerases are rather infrequent in absence of Tat.

### The DHS1 promoter nucleosome is infrequently detected in presence of Tat

SMF analysis shows a nucleosome footprint at the DHS1 region in ‘No Tat’ cells (Fig. 1D and 1F). This nucleosome covers the core promoter elements and in particular the TATA box and the SP1 binding sites, and it could thus contribute to the poor transcription observed in absence of Tat. This nucleosome further appears to be controlled by Tat because its presence decreases in ‘No Tat’ cells (Fig. 1). To investigate the role of this nucleosome in HIV-1 transcription, we first treated the ‘No Tat’ cells with the histone deacetylase inhibitor Trichostatin A (TSA), a known activator of latent viral transcription ^25,43,45^. Our reporter cell lines contained an intronic array of 128xMS2 stem loops and we thus assessed the effect of TSA treatment using live cell imaging with MCP-eGFP. We treated the ‘No Tat’ cells with 500 nM TSA and measured the intensities of HIV-1 transcription sites for 12h. We observed a progressive increase of viral transcriptional activity over the time course (Fig. 5A), while the untreated cells maintained a constant transcriptional activity (see below Figure 7A). Next, we treated ‘No Tat’ cells with 500 nM TSA for 8h and analyzed the molecular states of the promoter using SMF, using untreated ‘No Tat’ cells as a control. The data showed that the overall accessibility of the HIV-1 promoter was increased following TSA treatment (Fig. 5B). In particular, the binding probability of the DHS1 nucleosome decreased from 0.6 in untreated cells to 0.4 after TSA treatment (Fig. 5C and 5D). Conversely, the binding of TBP and RNAPII increased 2 to 4 fold (Fig. 5C and 5D). This was consistent with an increased accessibility of the promoter following TSA treatment, notably by displacing the DHS1 nucleosome that covers the core promoter elements.

**Figure 5:**
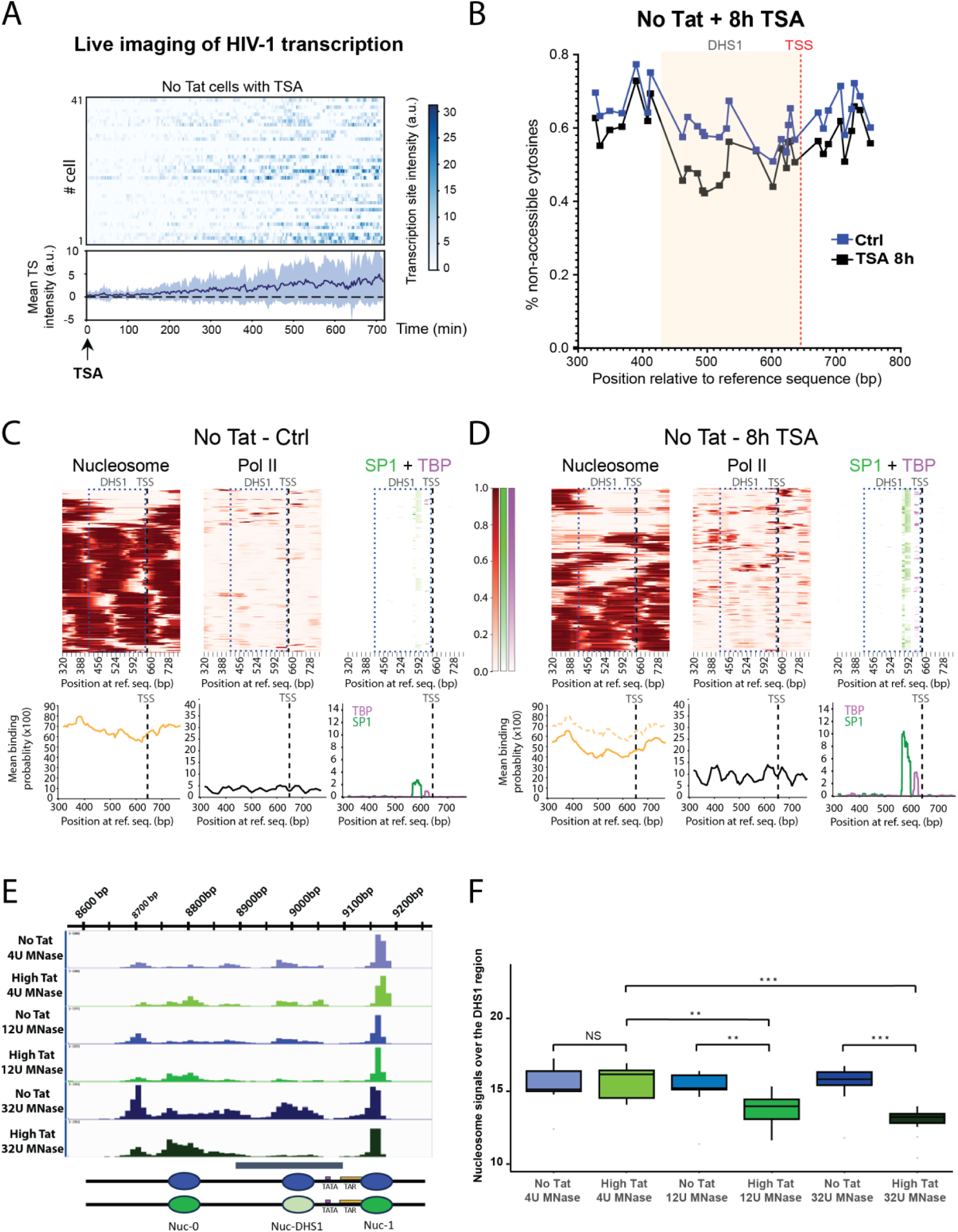
The DHS-1 region is occupied by a nucleosome in absence of Tat. **(A)** Live cell imaging of HIV-1 transcription in No Tat cells treated with 500 nM TSA over 8 h, with one image stack recorded every 3 min. The x-axis represents the time (in min) and each line of is a single cell. The intensity of the HIV-1 transcription sites is color-coded and the color scale is indicated on the right (in arbitrary units). Bottom panels: mean intensity of transcription sites. **(B)** Average accessibility of the GpC dinucleotides of the HIV-1 promoter in No Tat cells treated with TSA (black), as compared to non-treated control (blue). The yellow area marks the DHS1 region and the red dashed line marks the TSS. **(C-D)** Binding probabilities of nucleosomes, RNAP II and various transcription factors in No Tat cells treated with TSA **(D)**, as compared to a non-treated control **(C)**. Probabilities are estimated by Hiddenfoot. Top panels: each lane is a single molecule, blue rectangle gives the DHS1 area, the black dashed line marks TSS, and the color scale bars give the binding probabilities. Bottom panels: average binding probabilities for the indicated factors. SP1 is in green and TBP in purple. The dashed lines show the mean binding probabilities of nucleosome and RNAP II in untreated No Tat cells. **(E)** Read counts obtained with a MNase capture-Seq experiment in High Tat and No Tat cells, using increasing MNase concentrations. The area shown corresponds to the HIV-1 promoter, with the position of the nucleosomes indicates in the bottom schematic. **(F)** Quantification nucleosome signals over the DHS1 region, in High Tat and No Tat cells treated with increasing concentration of MNase. The signals correspond to normalized read counts.

To unequivocally confirm the presence of a DHS1 nucleosome, we sought to capture this information using capture MNase-seq. ‘High Tat’ and ‘No Tat’ nuclei were treated with increasing concentration of MNase and mononucleosome fragments were enriched using a custom-designed array of biotinylated RNA probes covering the HIV-1 reporter (Fig. S5C and 5D). Sequencing of the purified DNA fragments showed a clear nucleosome occupancy of the Nuc-0 and Nuc-1 regions that are known to harbor stable nucleosomes. Interestingly, the signals of the DHS1 region decreased with increased MNase concentration in the ‘High Tat’ cells, while it was stable in the ‘No Tat’ cells (Fig. 5E-F). This confirmed that the DHS1 region of the HIV-1 promoter harbors a nucleosome that is displaced in presence of Tat.

### Tat remodels the HIV-1 promoter chromatin by binding to the TAR RNA element

It is well established that Tat is recruited to the HIV-1 promoter by the nascent TAR RNAs. Because the disappearance of the DHS1 nucleosome is dependent on Tat, we investigated whether it also required the binding of Tat to nascent TAR RNAs. To this end, we introduced point mutations into the TAR element that prevented its binding to Tat and Cyclin T1 (Fig. 6A, S6A; ^58^). The resulting reporter, named ‘TAR-Double Mutant’ (‘TAR-DM’), was first assessed for its basal and Tat-induced activity using luciferase assays, showing that the activation by Tat was nearly abolished in the ‘TAR-DM’ promoter (Fig. S6B). We then introduced the mutant reporter into HeLa cells at the same locus as in the previous cell lines, generating ‘High Tat TAR-DM’ and ‘No Tat TAR-DM’ cells, which respectively expressed and did not express Tat. Of note, the ‘High Tat TAR-DM’ cells expressed Tat at comparable levels as the original ‘High Tat’ cells (Fig. S6C). We then used smFISH to quantify the level of transcription in the new cell lines. As expected, transcriptional activity in the ‘High Tat TAR-DM’ was greatly diminished as compared to the ‘High Tat’ cells (Fig. 6C). Nevertheless, the reporter was slightly more expressed in the ‘High Tat TAR-DM’ cells as compared to the cells lacking Tat (2-3 fold), possibly due to a residual binding of Tat to the mutant TAR RNA (Fig. 6B, 6C).

**Figure 6:**
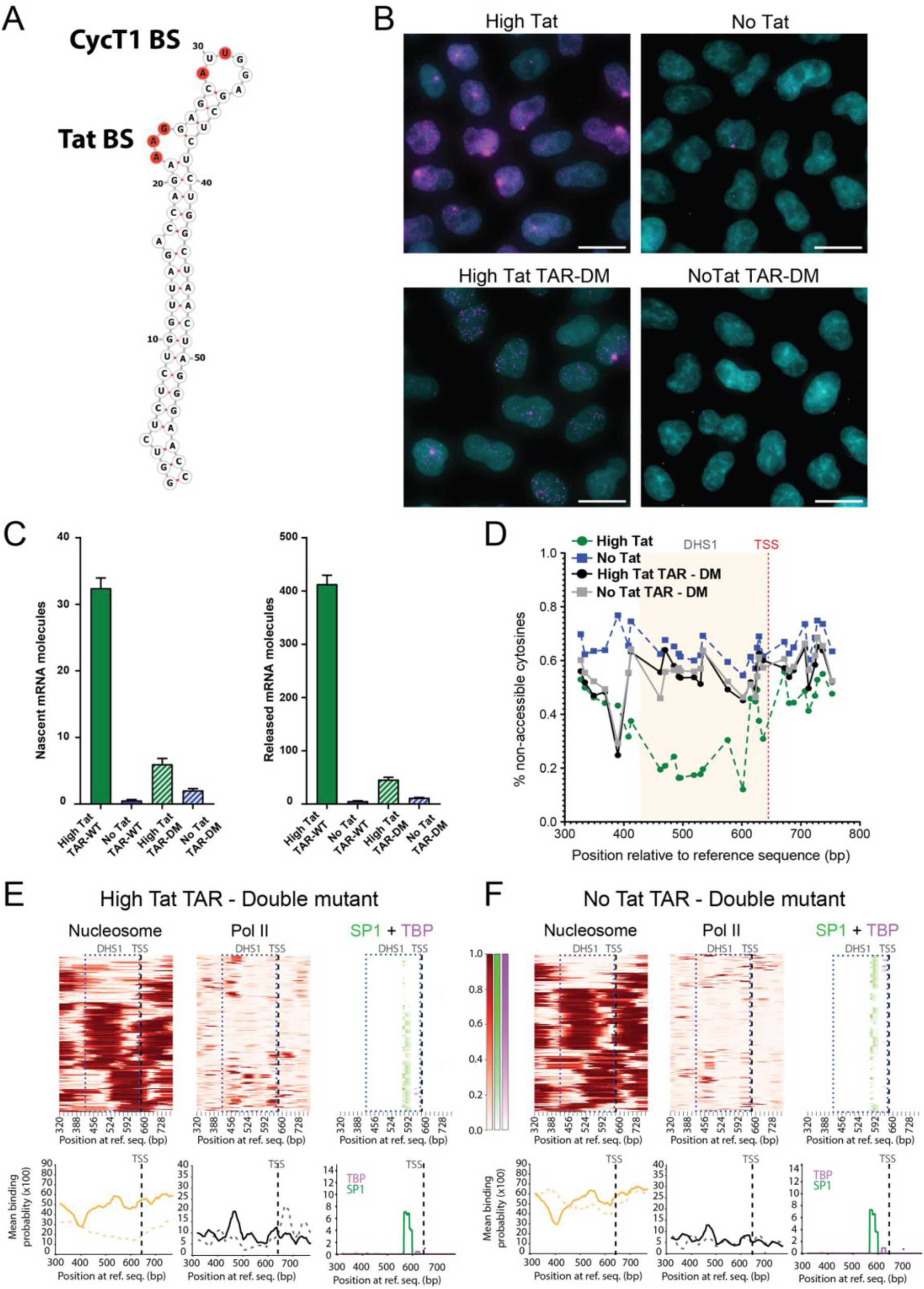
Tat remodels the HIV-1 chromatin via binding to the TAR RNA element. **(A)** Structure of TAR element. The nucleotides that have been modified to create the TAR double mutant are marked in red. **(B)** Images are micrographs of HeLa cells stably expressing the wild-type and TAR double mutant reporters, with (left) and without (right) Tat. Cells were labelled by smFISH with Cy3-labelled MS2 probes, and nuclei were stained with DAPI. Large bright spots are transcription sites and small dots are single molecules of RNA. Scale bars: 20 microns. **(C)** The bar plots show the quantification of nascent (within the transcription spot) and released nucleoplasmic RNA transcripts in High Tat and No Tat cells, with or without mutated TAR element. **(D)** Average accessibility of the GpC dinucleotides of the HIV-1 promoter in No Tat TAR-DM (grey) and High Tat TAR-DM (black) cells, as compared to No Tat and High Tat wild-type controls (blue and green dashed line, respectively). The yellow area marks the DHS1 region and the red dashed line marks the TSS. **(E-F)** Binding probabilities of nucleosomes, RNAP II and various transcription factors in High Tat TAR-DM **(E)** and No Tat TAR-DM **(F)** cells. Probabilities are estimated by Hiddenfoot. Top panels: each lane is a single molecule, blue rectangles gives the DHS1 area, the black dashed line marks TSS, and the color scale bars give the binding probability. Bottom panels: average binding probabilities for the indicated factors. SP1 is in green and TBP in purple. Dashed lines show the mean binding probabilities of nucleosome and RNAP II in High Tat and No Tat wild-type controls, respectively.

We then performed SMF to evaluate the molecular states of the promoter in the ‘High Tat TAR-DM’ and ‘No Tat TAR-DM’ cells. The bulk methylation showed nearly identical profiles in the two cell lines, showing that Tat had little effect on the HIV-1 promoter when it was unable to bind TAR. The profiles also displayed a striking resemblance to the one observed in the wild-type ‘No Tat’ cells, although some differences were seen in the Nuc-0 region, 200 nucleotides upstream the TSS (Fig. 6D). Most importantly, single molecule analysis showed that a nucleosome was frequently detected in the DHS1 region in the ‘High Tat TAR-DM’ mutant cells (average binding probability of 0.55). This was similar to the results observed in absence of Tat for both the wild-type and mutant TAR cell lines (Fig. 6E and 6F). This contrasted to the low binding probability observed in wild-type ‘High Tat’ cells (0.15; Fig. 1E). The increased nucleosome detection and lower transcription rate of the ‘High Tat TAR-DM’ cells was mirrored by the lower detection of TBP and RNAPII (Fig. 6E). This indicated that the TAR mutations abolished the effects of Tat and that Tat needed to bind to TAR to regulate the DHS1 nucleosome of the viral promoter.

### Combining live cell imaging and SMF for in-depth analysis of promoter states

Using live cell imaging of nascent RNAs, the signals of transcription sites can be deconvolved to extract the timing of individual polymerase initiation events ^48,59^. This yields the survival function, which is the distribution of the waiting times between successive initiation events. Different kinetic promoter models can be fitted to this survival function, yielding the transition rates between promoter states ^48,59^. However, these phenomenological models lack molecular interpretability of their states, which are usually simply referred to as ‘ON’, ‘OFF1’, ‘OFF2, etc. Conversely, SMF is a powerful tool to describe the molecular diversity of promoter states, but it only provides static snapshots. To construct kinetic models in which promoter states have well-defined molecular representations, SMF data need to be integrated with real-time measurements of transcription. To do so, we proceeded in three steps. First, we used transcription imaging data to estimate the survival function of initiations events. Second, we classified the SMF data into promoter states that are well defined molecularly, selected to reflect key steps of the initiation process (nucleosome removal, PIC assembly, polymerase loading and firing). Finally, we explored mechanistic hypotheses by finding the models for which the transition rates between the promoter states could simultaneously fit the survival function of transcription initiation and the state frequencies measured by SMF. Note that this imposes hard constraints because for a given model, the survival function also determines the frequencies of promoter states ^60^.

We had already imaged ‘High Tat’ and ‘No Tat’ cells ^6,48^. Yet, because of technical limitations, previous analyses used a combination of 8h long movies at a low temporal resolution, with 30 minutes movies at a high temporal resolution. Thanks to improvements in microscopy and in reliability of the deconvolution pipeline ^59^, we could now image ‘High Tat’ and ‘No Tat’ cells using a single movie of 12h with a resolution of one image per minute, while keeping a sensitivity of single polymerase throughout the movies. This removed errors caused by the stitching of the survival functions derived from short and long movies, a limitation of the former method ^59^. Using single nucleoplasmic RNAs from the live movies, we calibrated the signals of transcription sites and quantified the exact number of nascent RNAs over time in hundreds of cells. As expected, the ‘No Tat’ cells yielded only occasional bursts of transcription, with a mean of 0.4 nascent RNA per transcription site, while the ‘High Tat’ cells were nearly continuously active albeit with large variations, with a mean of 34 nascent RNAs per transcription sites (Figure 7A). We then deconvolved the movies and obtained the empirical survival function describing the distribution of time intervals between successive initiation events (see below Fig. S8B-C).

**Figure 7:**
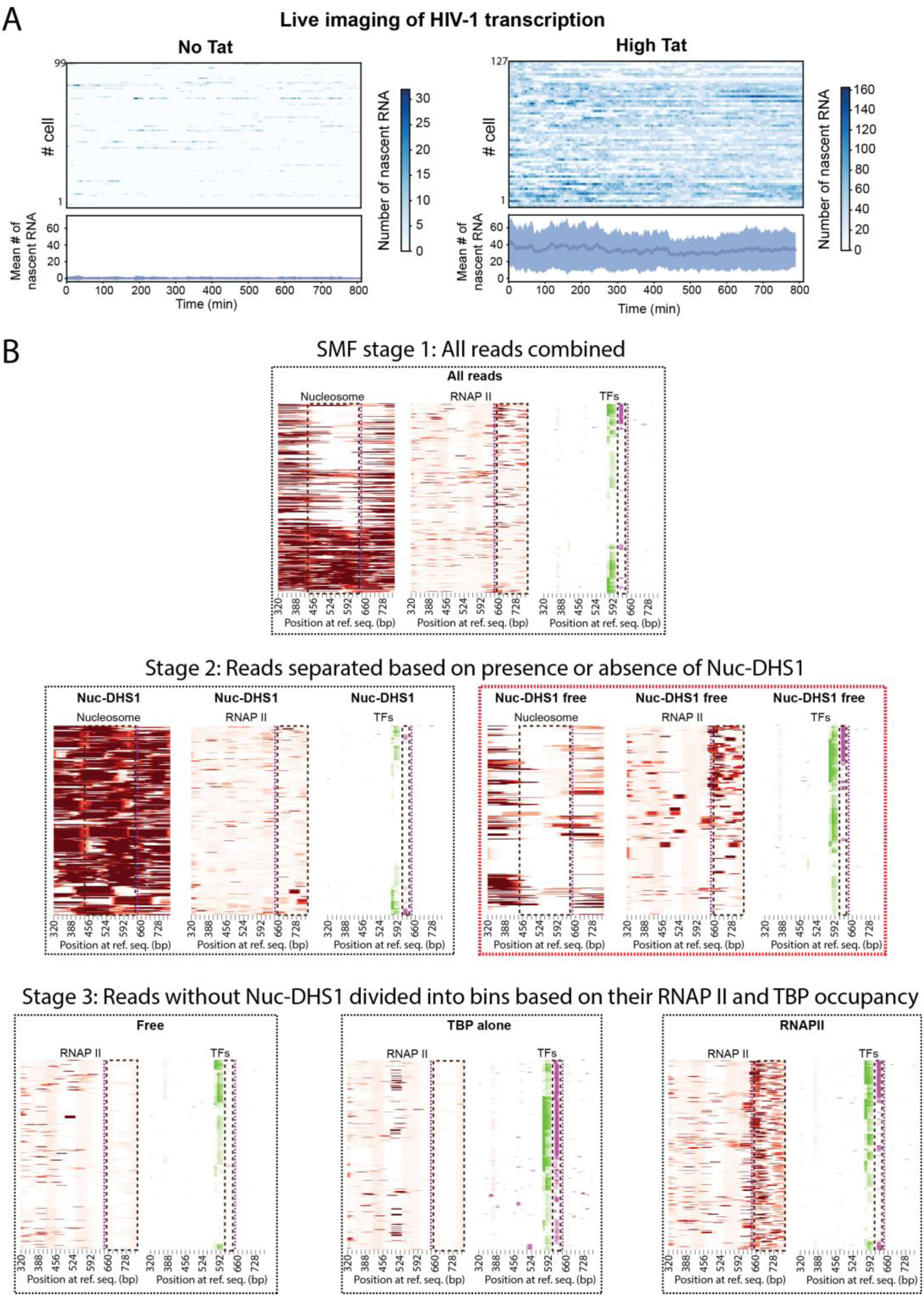
Combination of live cell imaging and SMF allows in-depth analysis of promoter states. **(A)** Live cell imaging of HIV-1 transcription in No tat (left) and High Tat (right) cells, with one image stack recorded every 3 min. The x-axis represents the time (in min) and each line is a single cell. The intensity of the HIV-1 transcription sites is color-coded and the color scale is indicated on the right (the unit is the number of nascent RNA molecules). Bottom panels: mean intensity of transcription sites. **(B)** Classification of single promoter molecules into four different classes based on their nucleosome, TBP and RNAPII occupancy. The molecular occupancy for each factor was determined by thresholding the single molecule binding probabilities given by Hiddenfoot as indicated in Fig. S2. All the SMF experiments were combined together (top panels; stage 1) and the SMF reads (corresponding to single promoters) were first divided into two groups depending on the presence or absence of a DHS1 nucleosome (middle panels; stage 2). SMF reads without a DHS1 nucleosome were subsequently divided into three additional groups (bottom panels): the first group contained neither TBP nor RNAP II (left, ‘free’ state), second group was bound by TBP only (middle, ‘TBP’ state), and the third group was bound by RNAPII (‘RNAPII’ state).

Next, we classified the binding states inferred from the SMF data into coarse-grained mesoscopic promoter states. First, we divided promoters into two classes, with and without the DHS1 nucleosome. The distribution of the DHS1 nucleosome binding probabilities was bimodal and we used k-means to optimally separate the two promoter classes (Fig. S2G). We then generated heatmaps showing their single-molecule occupancy profiles. The promoters with a DHS1 nucleosome (‘Nuc-DHS1’ state) were completely covered with nucleosomes including upstream and downstream of DHS1, indicating that this represents an inactive promoter class (Fig. 7B). The promoters without a DHS1 nucleosome were more diverse, with variable occupancies of Nuc-0, Nuc-1, SP1, TBP and RNA polymerases. We thus sub-classified the promoters devoid of a DHS1 nucleosome and made three additional classes (Fig. 7B; see Methods): (i) promoters also devoid of TBP and initiating polymerases (‘Free’ state); (ii) promoters having TBP but no initiating polymerases (‘TBP’ state); (iii) promoters having an initiating polymerase (‘RNAPII’ state). The advantage of this classification is that it can be ordered in a linear pathway easily integrated into a kinetic promoter model, starting with promoters having a DHS1 nucleosome, followed by the removal of DHS1 nucleosome, then TBP binding, and finally the presence of an initiating polymerase (see model in Fig. 8A). Using the thresholds of binding probability defined previously to determine the molecular occupancies at the levels of single molecules (see Fig. S2D, E, F), we quantified the fraction of molecules in each of the promoter classes, for the ‘High Tat’ and ‘No Tat’ cells (Fig. S7 and Table S1). These fractions reflected the frequencies of each promoter state in the overall cell population.

**Figure 8:**
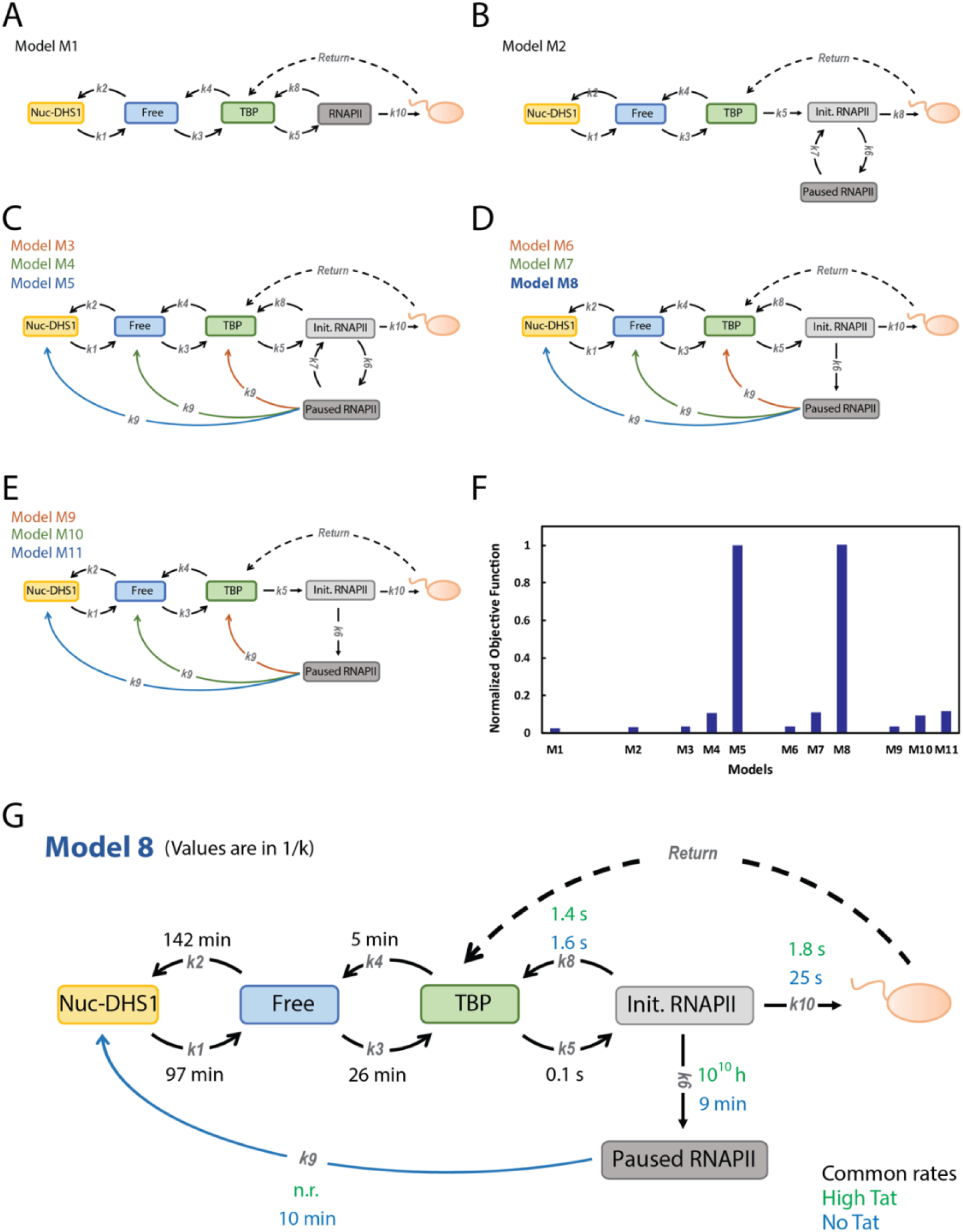
Integrative modelling of single molecule DNA footprinting with transcriptional imaging reveals key dynamics properties of the HIV-1 promoter. **(A-E)** Kinetic models describing the dynamic of the HIV-1 promoter. *k1-10* are different kinetics rates, orange balls are elongating RNAP II. Each model assumes that after elongation starts, the promoter immediately returns to the TBP state (dashed arrow). **(F)** Goodness of optimal solutions for each of the models. The values are the inverse of the objective function for the best parameters, normalized to that of the best model. **(G)** The best parameters for the winning model M8. The lifetime of transitions (1/k) are in minutes or seconds as indicated, and are provided for High Tat (green) and No Tat (blue) cells. n.r.: not relevant.

### Hybrid SMF/Live cell modeling reveals how the HIV-1 promoter functions and the mechanism of Tat regulation

Next, we aimed to fit a kinetic promoter model to the survival function derived from the MS2 transcriptional imaging data, using the frequencies of promoter states measured by SMF as constraints. To ensure realistic fits, we added two more constraints, one on the mean RNA production rate, which should match the values measured experimentally by smFISH; and the other to ensure identical promoter transition rates for ‘High Tat’ and ‘No Tat’ cells, for promoter states devoid of a polymerase. Indeed, Tat only acts on the HIV-1 promoter when a nascent RNA is present, i.e. on promoter states having a polymerase (see above Fig. 6). Overall, the objective function used during the fit (see Methods and Supplemental Note), simultaneously minimized the differences between the model and: (i) the two empirical survival functions of initiation events in ‘High Tat’ and ‘No Tat’ cells, as obtained from live transcription microscopy data; (ii) the frequencies of promoter states measured by SMF in the same two conditions, and (iii) the average RNA production rates measured by smFISH in the same two cell lines.

We tested multiple models to explore the mechanisms of transcription initiation. The simplest model that could account for the SMF data is a four states reversible linear model, going from a promoter with the DHS1 region occupied by a nucleosome (state ‘DHS1-Nuc’), to DHS1 free (state ‘Free’), TBP-bound (state ‘TBP’), and finally, a promoter with an initiating polymerase (state ‘RNAPII’), which can either elongate or abort. This model, termed M1 (Figure 8A), did not fit the data well. It resulted not only in high values of the objective function but was also unable to recapitulate the promoter state frequencies measured by SMF and the average RNA production rate (Fig. 8 and S8; Table S3). We thus sought more complex models. It is well established that the transition from initiating to elongating polymerases is a point of regulation that involves many modifications of the polymerase, including the phosphorylation of its CTD, the dissociation of negative elongation factors and the association of stimulating factors. Moreover, we previously observed that polymerases initiating on the HIV-1 promoter can sometimes divert from the productive pathway to enter long pauses ^48^. We thus hypothesized that the initiating polymerase can exist in two states, a productive state that can directly enter elongation (‘initiating RNAPII’), and an unproductive state (i.e. ‘paused RNAPII’) that must convert into the productive state before entering elongation (Fig. 8B). The corresponding model, termed M2, did also not fit the entire data (Fig. 8 and S8; Table S3). However, an improvement of the fit was obtained when considering similar models with an additional long cycle, in which the unproductive paused polymerase can be removed, thereby converting the promoter into one of the states lacking a polymerase (Fig. 8). We tested all the possible combinations of such long cycles, in which removing the unproductive paused polymerase can either yield a state with a DHS1 nucleosome, a nucleosome-free region or a bound TBP (Fig. 8C). Variants of these models were also tested, in which various steps of the long cycle were made irreversible. This generated models with long cycles that became irreversible from the ‘Paused RNAPII’ (Fig. 8C), the ‘Initiating RNAPII’ state (Fig. 8D), or the ‘TBP-bound’ state (Fig. 8E). A total of 9 models were obtained, out of which models M5 and M8 produced the best fits, well separated from all the other models (Fig. 8F). Models M5 and M8 have both a long irreversible cycle linking the ‘Paused RNAPII’ state to the ‘DHS1-Nuc’ state. The difference between them lies in the transition between the ‘Initiating RNAPII’ and ‘Paused RNAPII’, which is reversible for model M5 and irreversible for model M8.

The results were robust and showed no overfitting. Moreover, the transition rates between the states ‘DHS1-Nuc’, ‘Free’, and ‘TBP’ were identical for model M5 and M8, further indicating their robustness. The parametric uncertainty was also small for both models, with few exceptions that could be explained (Table S3). For instance, the rate k8 was uncertain in ‘High Tat’ cells for both models M5 and M8. However, k8 is the rate of polymerase abortion and in ‘High Tat’ cells, this rate can be neglected as polymerase reloading is very fast and the long cycle k6-k9 is never used. Model M5 showed uncertainties for k9 and k6 in ‘High Tat’ condition, and for k7 in ‘No Tat’ cells. This model is more complex as the ‘Initiating RNAPII’ / ‘Paused RNAPII’ transition is reversible and the corresponding k6 and k7 rates can likely compensate each other to achieve the same effects.

A comparison of the fit results in ‘High Tat’ and ‘No Tat’ reveals the effects of this key viral activator. The promoter is very active in presence of Tat, as it continuously moves from the ‘TBP’ to the ‘initiating RNAPII’ state and then back to the ‘TBP’ state when the polymerase enters elongation. This behavior breaks in absence of Tat, because some polymerases enter an unproductive state, the ‘Paused RNAPII’ state, which leads to the removal of the polymerase and the occupancy of the DHS1 nucleosome, effectively closing the promoter. Preventing polymerase to enter the unproductive pathway is at the basis of the effect of Tat. In No Tat cells, the irreversible cycle is often used (once every 20 polymerases), leading to promoter closure and the shut-down of transcription for long time periods (∼1.5 hours). In contrast, this never happens in the High Tat cells, leading to high and sustained transcription levels, ∼100 fold higher than in absence of Tat. The key point of regulation thus occurs when the promoter loads an initiating polymerase. Indeed, this polymerase faces two irreversible competitive steps, to either move into elongation or enter an unproductive state ultimately leading to transcriptional shut-down. This competition has the hallmarks of a kinetic proofreading mechanism, which Tat largely shifts in favor of the productive pathway.

## Discussion

Imaging transcriptional activity in live cells has shown that promoters are highly dynamic and fluctuate between active and inactive states on multiple time scales, from minutes to hours ^6,7^. These transitions are essential as they are key point of regulation and their rates determine the overall promoter output. However, characterizing promoter dynamics in molecular terms has proven challenging ^1,3,4,11,12^. Here, we overcame this critical bottleneck by combining live cell transcription imaging with single molecule DNA footprinting (SMF). These methods provide highly complementary datasets: live imaging enables a detailed kinetic characterization of promoter activity ^59^, while SMF determines and quantifies the molecular diversity of promoter states ^10^. As such, transition rates can be inferred for each of the molecular states of the promoter. By analyzing the HIV-1 promoter in presence and absence of Tat, this approach estimates the kinetics of key molecular steps regulating viral transcription: nucleosome deposition/removal, PIC assembly, polymerase loading and pause release. It further explains how Tat indirectly controls the promoter chromatin by boosting elongation, and it reveals a crucial kinetic proofreading step that is alleviated by Tat.

### The DHS1 promoter nucleosome is indirectly regulated by Tat

The DHS1 region of the HIV-1 promoter is known to be more accessible than the Nuc-0 and Nuc-1 regions that are located immediately upstream and downstream. While generally considered nucleosome-free, a labile DHS1 nucleosome has been described in some studies ^47,61,62^. Our SMF and MNase data are consistent with these results and further highlight the importance of this nucleosome. Indeed, it not only covers key promoter elements such as the SP1 binding sites and the TATA box, but it also appears to act as a trigger for transcription: promoters having a DHS1 nucleosome are completely covered by nucleosomes, including upstream and downstream of DHS1, while promoters without a DHS1 nucleosome occur in a variety of states, with and without Nuc-0 and Nuc-1 nucleosomes, TBP or initiating polymerases. Displacement of the DHS1 nucleosome is thus a key step to activate the viral promoter. Importantly, our results indicate that Tat controls the DHS1 nucleosome but does so indirectly, by promoting elongation and preventing the formation of immobilized polymerases. The equilibrium between the rates of loading and removal of the DHS1 nucleosome should yield a frequent nucleosome occupancy of the DHS1 region (∼40%; see k1 and k2 in Fig. 8G). However, in the presence of Tat, the promoter constantly cycles between TBP-occupied and polymerase-loaded states, driving the equilibrium toward open chromatin states. This cycle breaks in absence of Tat because paused polymerases frequently fail to enter elongation, leading to promoter closure and nucleosome occupancy of the DHS1 region. The chromatin of the core promoter is thus indirectly regulated by Tat, without a direct alteration of the rates of nucleosome deposition or removal.

Several studies have used inducible histone expression systems and dynamic ChIP to measure the turn-over of nucleosomes over entire genomes ^63–66^. This has revealed that, in general, nucleosome turn-over is rapid at enhancers and promoters. In mES cells, the dynamics of the nucleosomes localized immediately upstream TSSs, at a site similar to the DHS1 region in the HIV-1 promoter, weakly anti-correlates with occupancy ^63,65^. In yeast, the dynamics of promoter nucleosomes have been estimated to be in the range of 0.6 -1 exchange per hour ^64^. This is remarkably similar to our findings for the DHS1 nucleosome on the HIV-1 promoter, for which we estimate a lifetime of ∼90 minutes and a loading rate of 0.5 per hour.

### PIC assembly

We find that once the DHS1 nucleosome has been removed, TBP is loaded at a relatively slow rate (one event per 25 minutes), while its removal is fast (once every 5 minutes). BTAF1 has been shown to remove TBP from chromatin in an ATP-dependent manner. Moreover, FRAP and single particle traking studies in mammalian cells have estimated that TBP is removed once per 1.5 minute on random chromatin location ^67,68^, with a loading rate ∼3 fold slower than its removal ^67^, similar to the on/off rate ratio that we estimate here for the HIV-1 promoter. Dynamic ChIP in human HEK293 cells has measured a turn-over rate of ∼30 minutes for TBP on promoters with a TATA box ^69^, and a correlation between high transcription activity and slow TBP turn-over. The HIV-1 promoter has a TATA box and transcribes highly in presence of Tat. In addition, its TBP/TATA box interaction was shown to control long time scales of promoter ON periods ^6^, indicating a long-lived interaction that promotes multiple rounds of initiation from a single TBP binding event. This points to a long residency time of TBP on the HIV-1 promoter when Tat is present. Our model indicates that once TBP is loaded on the TATA box, the rate of PIC assembly and polymerase loading is very fast. In the presence of Tat, polymerases are efficiently fired, leading to a constant cycling of the promoter between TBP-bound, initiating polymerase, and back. The model estimates that this cycle would last on average 1.5h, resulting in long residence times of TBP on the TATA box. Single protein tracking of PIC components and RNA polymerase II in yeast has shown that PIC assembly and polymerase loading is indeed a fast process occurring in a few seconds after TBP binds DNA ^70^. It also showed that the presence of polymerase on the promoter increases the residency time of the PIC, in agreement with our results.

### Kinetic proofreading of paused polymerases

Once loaded on the promoter, the initiating polymerase must undergo a series of phosphorylation and must change partners to enter productive elongation ^33,38^. Alternatively, initiating polymerases can also immediately abort and return the promoter to the TBP-bound state. Genome-wide data measuring RNAPII turn-over at promoters, or the tiny RNAs associated with initiation, suggests that abortive polymerases are rather frequent ^10,72–74^. Interestingly, the models M9-11 lack a direct abortive pathway (k8) and do not fit the data well, indicating that polymerase abortion is required to capture the entire dynamic of the HIV-1 promoter. Our data also indicate that while PIC assembly and polymerase loading is very fast (k5 ∼ 10 s^-1^), its escape from the promoter (k10) is slow in absence of Tat and becomes rate-limiting (k10 < k5). Consequently, most of the initiating polymerase immediately abort (84%; k8 ∼ 0.5 s^-1^). In contrast, promoter escape is ten times more rapid in presence of Tat (k10 ∼ 0.5 s^-1^), such that about half the polymerases immediately abort while the other half goes into productive elongation. This value consistent with recent live imaging of RNAPII on the HIV-1 promoter using the same reporter cell lines ^71^. A lower proportion of immediate abortion is also fully consistent with the function of Tat, which is to promote elongation by recruiting P-TEFb to the polymerases ^25–27^.

In addition to immediately aborting, initiating polymerases can also be removed via a long irreversible cycle, which links polymerases arrested in long pauses to the deposition of a DHS1 nucleosome. This is indeed a key feature of the hybrid MS2/SMF model because only models incorporating this long cycle can fit the experimental data, highlighting its importance. Initiating polymerases thus appear to face two irreversible competing steps, one leading to productive elongation and recycling of the promoter to a highly transcription-competent state (e.g. TBP-bound), and the other leading to a long pause, polymerase removal and DHS1 nucleosome deposition, thereby preventing new initiation events for long time periods (∼1.5h). This indicates a kinetic proofreading mechanism, in which initiating polymerases have a certain time to enter elongation (9 minutes in absence of Tat), and if they fail to do so, trigger the shut-down of transcription. This contributes to the low level of transcription observed in absence of Tat, as on average it happens once for every 20 polymerases that enter elongation, while it never happens when Tat is present. This points to a key role of P-TEFb in the proofreading mechanism, because P-TEFb is the key factor controlling the switch from initiation to elongation, and initiating polymerases lack P-TEFb in absence of Tat ^25–27^.

The benefits of irreversible steps and kinetic proofreading mechanism during transcription initiation have been previously discussed ^75^. The polymerase proofreading mechanism observed here for HIV-1 could be more general and may play an important role genome-wide. Vertebrate genomes are pervasively transcribed and pausing/early termination frequently discriminates functional from spurious initiation events ^76^. The proofreading mechanism described here could recognize defective polymerases that fail to enter elongation, abort them and close the chromatin to prevent new initiation events. A number of protein complexes have been identified that link chromatin remodelers with paused RNAPII, including some recognizing S5 and Y1 phosphorylated forms of the RNAPII CTD ^77–80^. Conceivably, an initiating polymerase phosphorylated on S5 of its CTD but failing to become phosphorylated on S2 by P-TEFb would get stuck on the promoter and may recruit these complexes to achieve long-term transcriptional shut-down, thereby preventing pervasive transcription. In agreement with this possibility, a lack of CTD S2 phosphorylation in *S. Pombe* has been shown to drive the occupancy of core promoters by nucleosomes, for hundreds of genes ^81^. The mechanism involves the direct recognition of the arrested S5P polymerases by the Set1-WD82 complex, which then methylates histones H3 on K4. This in turns enables the recruitment of histone deacetylase complexes such as SET3C and, ultimately, promoter closure. Future studies may address whether similar molecular mechanisms operate in mammalian cells and enable a kinetic proofreading of RNA polymerases.

## METHODS

### Cell lines and culture

Human HeLa Flp-in H9 cell lines were cultured in complete DMEM, supplemented with 10% FBS and penicillin/streptomycin (10U / ml). Cells were kept in a humidified atmosphere with 5% CO2 at 37 °C. Stable expression of MCP-eGFP and TetR-tagBFP driven from the ubiquitin promoter was accomplished by lentiviral infection. The MCP contained the deltaFG deletion and the V29I mutation. Both MCP-eGFP and TetR-tagBFP contained an SV40 NLS. Positive cells were selected by FACS. Only cells expressing low levels of fluorescence were taken. Stable cell lines expressing different versions of HIV-1 LTR reporter were created using the Flp-In system as described in ^6^. Briefly, cells were transfected with the MS2*128 HIV-1 reporter plasmids (all reporter plasmids are listed in Table S4) using JetPrime (Polyplus), following the manufacturer’s protocol. Positive clones were identified using Hygromycin selection (150 µg/ml) and analyzed by smFISH and genotyping.

The ‘No Tat’ cell line expressed the MS2*128 HIV-1 reporter gene but no Tat protein. To produce ‘High Tat’ cell lines, the Tat-Flag fusion protein under the control of chicken beta-actin promoter was stably introduced in cells by CRISPR-Cas9 using an AAVS1 repair vector. For the ‘High Tat SP1-mutant’ and ‘High Tat TATA-mutant’, the pSpoII-Tat plasmid was used instead of AAVS1 CRISPR. Here, the Tat-Flag cDNA is followed by an IRES-Neo selectable marker and is transcribed under the control of the CMV promoter. Positive clones were selected on puromycin at 2 μg/ml or neomycin at 400 μg/ml, respectively. Tat expression was verified by Western blot, while localization and expression homogeneity were confirmed by immunofluorescent staining using antibodies against Flag. Selected clones were regularly re-tested to ensure the expression stability of Tat between experiments.

### Antibodies and drug treatments

Western blots were performed with α-Flag antibody (Sigma F3165) at dilution 1:1000 and detection was done with fluorescently labeled α-mouse secondary antibody (Abcam ab6728) at 1:10000. Immunofluorescence was done using the same α-Flag antibody at 1:300 and α-mouse-Cy3 secondary antibody (Jackson 115-166-006, 1:1000). Cell treatments were performed with drugs at the following concentrations: Triptolide, 1mM (Sigma T3652), Trichostatin A, 500 nM (Selleckchem S1045).

### Plasmids and reporter constructs

Plasmid maps and sequences are available upon request. The 128xMS2 HIV-1 reporter and High Tat expression vector were described previously ^6,48^. The HIV-1 LTR TAR-MS2x4 constructs 4c2 and 3c3 were created by inserting four MS2 stem loops inside the TAR stem. Each MS2 stem-loop had a different sequence to prevent cross-hybridization between themselves. The stem-loops were separated from each other as well as from the core TAR loop by 3 nucleotides. Proper folding of corresponding RNAs was verified by mFold ^82^. The final sequences were synthesized by GenScript and sub-cloned in an HIV-1 reporter vector (pINTRO-FRT-Hgr-HIV-1-SacB*64), which contained an insertion of 64 SacB stem-loops in the intron instead of the original MS2*128 stem-loops, as well as an FRT-Hygro cassette for Flp-in recombination.

The HIV-1 TAR-DM mutant construct was created by site directed mutagenesis using Agilent’s QuikChange II-E Site-Directed Mutagenesis Kit (Agilent, 200555) following the manufacturer’s protocol and using the mutagenesis primers 5’ GTCTCTCTGGTTAGACCAGAaagGAGCCTGGGAGCTCTCTGGC and 5’ CTGGTTAGACCAGATCTGAGCaatGGAGCTCTCTGGCTAACTAG that target the Tat and Cyclin T1 binding sites, respectively, with the mutations indicated in lowercase. Successful introduction of the mutations was verified via Sanger sequencing and the construct with double mutation was then subcloned into the pINTRO-FRT-Hgr-HIV-1-MS2*128 reporter vector.

### Luciferase assays

To test the transcriptional activity of the HIV-1 LTRs, the core promoter within the LTR was subcloned into pGL3 (Addgene, #212936) in front of the firefly luciferase gene. The resulting vectors were transfected into the HeLa cells together with pSV-Tat or an empty vector (pBlueScript, Stratagene 212205) using JetPrime (Polyplus), following manufacturer’s protocol. The vector pTK-RL (Promega, E2241), which expressed renilla luciferase, was co-transfected as an internal control to correct transfection efficiency. Luciferase expression was measured on a BertholdTech TriStar LB 941 (on 96-well, measurement counting time 1 sec, no emission filter) using the Dual-Glo^TM^ Luciferase assay system (Promega, TM058).

### SmFISH / smiFISH and image quantification

Cells were grown on 12 mm round coverslips, fixed in 4 % formaldehyde/1xPBS for 20 min at RT and permeabilized in 70% ethanol at 4°C overnight or longer. Prior to the experiment the ethanol was rinsed with 1xPBS and samples were pre-incubated with 40% Formamide/1xSSC for 30 min before being hybridized with a mix of fluorescent DNA oligonucleotides as previously described ^57,83^. SmFISH oligonucleotides were directed against the MS2*32 and MS2*4 repeats and were directly conjugated with Cy3 (four molecules of Cy3 per oligo; ^57^), while the 64*SacB sequence was detected using smiFISH oligos ^83^, which were indirectly labelled with a Cy5-Flap-Y oligo. Hybridization was done overnight at 37 °C in a humidified chamber as previously described ^83^. Following hybridization, coverslips were washed twice 30 minutes at 37°C in 1xSSC, 15% formamide, once in PBS and mounted in Vectashield (VectorLabs).

Imaging was performed on a Zeiss Axioimager Z2 Apotome, equipped with a LED Xcite 120 LED illumination controlled by Zen and an ORCA-Flash4 LT camera (Hamamatsu). Acquisition was performed in 3D with a Z-spacing of 0.3 µM and a 63x Apochromat 1.4 NA oil objective.

SmFISH image analysis was performed using SmallFISH (available at: https://github.com/2Echoes/small_fish_gui), a custom-made user-friendly software based on the BigFish pipeline ^84^. For TAR-MS2*4c2 and TAR-MS2*4c3, the quantifications were done manually due to the low signal-to-noise ratio of the MS2*4 probes that prevented automated spot detection. We counted the cells with co-occurrence of TetO*96 (tagBFP) and MS2*4 (Cy3) signal, but lacking a SacB signal (Cy5). To estimate the smFISH background and the rate of false detection, we also counted the colocalization of the tagBFP and Cy3 signals in cells that lacked the HIV-1 reporters.The proportion of cells with a false positive MS2 signal (∼10%) was then deduced from the numbers obtained in the HIV-1 reporter cell lines, yielding the fraction of cells with a paused polymerase.

### Single-molecule DNA methylation footprinting

The HIV-1 reporter cell lines were seeded at a confluency of 7x10^5^ cells per 3.5 cm diameter Petri dish and left to grow overnight. Cells were then trypsinized, washed twice with ice cold PBS and counted. 250,000 cells per sample were lysed in ice cold lysis buffer (10 mM Tris-HCl pH 7.4, 10 mM NaCl, 3 mM MgCl2, 0.1 mM EDTA, 0.5% NP40) for 10 min on ice, followed by centrifugation at 3000g for 5 minutes. Nuclear pellets were washed with SMF wash buffer (10mM Tris pH 7.4, 10mM NaCl, 3mM Mgcl2, 0.1mM EDTA), spun as above and subsequently resuspended in 1x M.CviPI reaction buffer (50 mM Tris pH 8.5, 50 mM NaCl, 10 mM DTT). Nuclei were treated with 200 U of M.CviPI (NEB M0027L) at 37 °C for 7.5 min in presence of 0.6 mM SAM and 300 mM Sucrose. 100 U of M.CviPI and 128 pmol of SAM was subsequently added before a second round of incubation at 37 °C for 7.5 min. Reaction was stopped by addition of Stop buffer (20mM Tris pH 7,5, 600 mM NaCl, 1% SDS, 10mM EDTA) and incubated overnight at 55 °C. DNA was extracted by phenol-chloroform extraction followed by glycogen / isopropanol precipitation. DNA was then resuspended in water, treated with RNAse A for 30 min at 37 °C and quantified using the Qubit 1x dsDNA HS assay kit (ThermoFisher Scientific Q33231).

1500 ng of extracted DNA was used as an input for bisulfite conversion with the EpiTect Bisulfite Kit (Qiagen 59104) according to the manufacturer’s protocol. The HIV-1 core promoter as well as control regions (encompassing CTCF binding sites near the ABL2 and TP53 genes) were amplified using Kapa HiFi HotStart Uracil+ Ready Mix (Roche KK2802) with the following cycling conditions: denaturation at 95 °C for 3 min; denaturation, annealing and elongation for 34 cycles respectively at 95 °C, 55 °C and 72 °C for 20 s, 15 s and 1 min; final elongation at 72 °C for 1 min. The primers were designed using the Primer 3 software to hybridize to the *in silico* bisulfite converted template and are listed in Table S5. The amplicons (400 to 500 bp) were pooled together and purified using AMPureXP magnetic beads (Beckman Coulter A63880). Libraries were prepared using NEBNext Ultra II DNA library prep kit for Illumina (NEB E7645) and were evaluated on a Bioanalyzer 2100 using Agilent DNA 1000 kit (Agilent 5067 – 1504). Pools of up to 12 libraries were sequenced on a Miseq instrument using 300 bp paired-end sequencing.

Data were analysed using the SingleMoleculeFootprinting Bioconductor package as previously described ^10,49^. Briefly, reads were trimmed to minimal length of 249 bp using Trimmomatic (https://doi.org/10.1093/bioinformatics/btu170) with the following settings: “ILLUMINACLIP:TruSeq3-PE.fa:2:30:10 LEADING:3 TRAILING:3 SLIDINGWINDOW:4:15 MINLEN:249” to remove adapters and Illumina specific sequences as well as all incomplete reads. Trimmed sequences were aligned to bisulfite converted human genome (BSgenome.Hsapiens.UCSC.hg38, available as Bioconductor package) and the HIV-1 LTR sequence (custom forged BSgenome, available upon request). Bulk methylation as well as single molecule methylation results were obtained using the MethCall function of the SingleMoleculeFootprinting package.

The HIV-1 specific primers hybridized to a region that is common for both 5’ and 3’ HIV-1 LTR. However, in our construct the 3’ LTR contains nine SNPs that are not present in 5’LTR. We took advantage of this and sorted out the reads that were specific for the 5’ LTR specific using a custom script written in Python (available upon request). Finally, all the reads containing NA at any position were removed before plotting and subsequent single molecule processing.

### Analysis of SMF using HiddenFoot

We used Hiddenfoot to infer binding probability maps of nucleosomes, TFs, and RNAPII at the level of single promoters. This is a computational framework that integrates SMF data with a biophysical model of protein-DNA interactions ^52^. Briefly, HiddenFoot employs a thermodynamic approach based on Gibbs-Boltzmann statistics, systematically evaluating all possible non-overlapping binding configurations of TFs, RNAPII, and nucleosomes for each sequenced DNA molecule. Each configuration is assigned a statistical weight according to its sequence-specific binding energies and its consistency with the observed methylation pattern, under the assumption that protein binding prevents methylation. Binding affinities for TFs are computed using known TF binding motifs and their associated position weight matrices (PWMs), while RNAPII and nucleosome binding are modeled as sequence-independent. Finally, HiddenFoot assumes that TFs, when bound, protect DNA segments equal in length to their respective PWMs, while RNAPII and nucleosomes protect predefined lengths set by the user.

HiddenFoot incorporates several key parameters: the probabilities q_U_ and q_B_, which represent the probability of a cytosine being unmethylated when unbound or bound by a protein, respectively, accounting for the experimental efficiency of the SMF protocol; the local concentrations of TFs, RNAPII, and nucleosomes, which modulate their effective binding propensities; and, the inverse temperature parameter (β), which controls the model’s sensitivity to differences in binding energies. HiddenFoot employs dynamic programming to efficiently sum over all possible configurations, enabling parameter inference via stochastic gradient descent to maximize the likelihood of the observed methylation patterns. Finally, HiddenFoot outputs probabilistic occupancy profiles, representing the likelihood that each nucleotide is bound by a TF, RNAPII, or a nucleosome.

For the analysis of the HIV-1 promoter, we ran HiddenFoot using 15 PWMs from the JASPAR database (except for SP1 that was obtain from SwissRegulon), corresponding to TFs with high-affinity sites present within the promoter region (see Fig. 1C), and used protection lengths of 147 bp for nucleosomes and 40 bp for RNAPII. HiddenFoot running options were set to the default values and model parameters were optimized independently for each experimental condition.

### Classification of single reads into molecularly interpretable promoter states

To classify single DNA molecules into biologically meaningful promoter states, we pooled together the binding profiles inferred from Hiddenfoot from all the experimental conditions. We divided nucleosome occupancy into three distinct regions: Nuc-0, covering the upstream 150 bp; Nuc-DHS1, located at the transcription start site (TSS) and overlapping the TATA box and DNase hypersensitive site (DHS1); and Nuc-1, covering the downstream 150 bp and overlapping the elongation zone of RNAPII. Similarly, RNAPII binding was measured over the region downstream the TSS, and TBP binding over the TATA box.

For each DNA molecule, the mean binding probability across each region was computed, resulting in a simplified molecular profile capturing the key chromatin and transcriptional states. These profiles were concatenated across experimental conditions and hierarchically clustered to group molecules based on their regulatory state. We first applied k-means clustering to separate molecules into two categories based on Nuc-DHS1 occupancy: one group where the DHS1 region was nucleosome-occupied (representing a closed promoter) and another where the DHS1 region was nucleosome-free (an open promoter). Molecules classified as having an open promoter were further subdivided according to TBP and RNAPII binding. We classified molecules as belonging to one of 3 sub-states: free promoter (neither TBP nor RNAPII bound), TBP-only (bound by TBP but not RNAPII), RNAPII (bound by RNAPII). To this end, empirical thresholds were applied to define binding events at the single-molecule level. RNAPII was considered bound above a binding probability of 0.05, while TBP was considered bound above a binding probability of 0.005. These thresholds were selected on the basis of their ability to minimize false positives, using Triptolide and TATA mutant cells as controls (see Fig. S2). The fraction of molecules in each of these states was quantified for each condition and replicates were used to estimate experimental variability and generate error bars for each of the promoter states.

### MNase capture-seq

‘High Tat’ and ‘No Tat’ cells were seeded at a confluence of 2 x10^6^ cells per 10 cm diameter Petri dish and left to grow overnight. Cells were then trypsinized, washed twice with ice cold PBS and lysed with hypotonic buffer (10mM Tris HCl pH 7.5, 10 mM NaCl, 3 mM MgCl_2_, 0.5 % Triton X-100) for 15 min on ice, followed by centrifugation for 15 minutes at 500g. Nuclear pellets were washed with MNase digestion buffer (10mM Tris-HCl pH 7.5, 15 mM NaCl, 60 mM KCl, 3 mM CaCl_2_), spun and resuspended again in MNase digestion buffer. After initial preincubation (2 min at 37 °C), nuclei were treated with 4U, 12U and 32U of MNase (ThermoFisher Scientific 10848990) for 4 min at 37 °C. Reactions were stopped by addition of Stop buffer (20 mM EDTA, 20 mM EGTA, 0.4 % SDS and 0.5 mg/ml Proteinase K) and incubated overnight at 55°C. DNA was extracted by phenol-chloroform extraction followed by ethanol precipitation. DNA was separated with a 2 % agarose gel and mononucleosome fraction was cut out and purified using NucleoSpin Gel and PCR Clean-up kit (Macherey-Nagel, 740609). To obtain non-treated control, DNA was extracted from nuclei using Wizard Genomic DNA Purification kit (Promega, A1120) and sonicated with Bioruptor Pico (Diagenode) to produce DNA fragments 100-300 bp on average. Sonicated DNA of pINTRO-MS2x128 plasmid was also included as an independent control.

Libraries were prepared using SureSelect XT HS2 DNA reagent kit with indexes (Agilent G9981A) according to manufacturer’s protocol. 150 ng of input DNA was taken for each library (except for plasmid DNA where only 150 pg were used). To enrich for the HIV-1 reporter sequences, we used an array of biotinylated RNAs that were custom-designed by Agilent Technologies (Agilent 5191-6925) and that hybridized to the whole HIV-1 reporter vector except for the sequence of the MS2x128 stem loops. Bait capture was performed using SureSelect XT HS2 Fast-Hyb Target Enrichment Kit (Agilent G9987A) following manufacturer’s guidelines. Libraries were pooled together prior to bait hybridization to reach a total of 1500 ng per pool. The library pool was concentrated using a vacuum concentrator (< 45 °C) and reconstituted with water (total volume 12 µl). Pooled DNA was hybridized with capture probes followed by an overnight hold at 21 °C. Hybridized DNA was captured using MyOne Streptavidin T1 Dynabeads (Thermo Scientific 10531265) and post-capture PCR was performed on bead-bound DNA via biotinylated-RNAs (16 cycles, SureSelect Post-Capture Mix). Post-capture libraries were purified using AMPure XP beads (Beckman Coulter A63880) and assessed by Bioanalyzer 2100 using Agilent HS DNA kit (Agilent 5067 – 4626). Capture MNase-seq libraries were sequenced on Novaseq with paired-end 50 bp settings.

### Processing and analysis of MNase-seq data

Paired-end Mnase-seq reads were trimmed of adapter sequences using Trim Galore tool (Trim Galore 0.6.6), combining Cutadapt (Cutadapt 1.18) with a quality control step (FastQC v0.11.9). Clean reads were aligned on the viral HIV DNA pNL4-3 sequence (GenBank: AF324493.2) using BurrowsWheeler Aligner (BWA 0.7.17). The aligned reads were filtered and sorted using Samtools 1.14. Duplicated reads (e.g. due to PCR) were removed using picard tools (picard-2.26.4; https://broadinstitute.github.io/picard/). For specific detection of nucleosome signal, coverages were obtained using Deeptools bamCoverage (deeptools-3.5.2) by filtering fragment of sizes between 100 bp and 200 bp with the ‘--Mnase – extendReads’ options to get the signal around the site of the nucleosome center. Genome browser visualization was performed on IGV using a custom reference genome created from pNL4-3 sequence (GenBank: AF324493.2). Boxplots were done with R 4.1.3 version and ggplot2.

### Live cell imaging

For live imaging of ‘High Tat’ and ‘No Tat’ cell lines, cells were seeded into 35 mm diameter Fluorodish (World Precision Inst. WPI, WFD35-100) and imaged in non-fluorescent media (Gibco^TM^ FluoroBrite^TM^ DMEM A1896701 complemented with 10% FCS) with a Spinning Disk Dragonfly microscope (Andor Technology), equipped with a 488nm illumination laser; an EMCCD camera iXon888 Life Andor 1024*1024 with a pixel size of 13 µm; and GFP emission filter 525/50. Acquisition was performed in 3D with a Z-spacing of 0.6 µM, 21 slices in total and a frame rate of one 3D stack per minute, using a 100X Plan Apo lambda 1.45 NA 0.13 mm WD oil objective. Cells were kept in a temperature-controlled chamber at 37 °C with 5% CO_2_ for the entire movies (13h).

### Movie analysis

Movie analysis was performed with a custom-made python software named bigFishLive, based on the bigFish analysis pipeline ^84^. Briefly, for each field of view, the nuclei were cropped and centered using an automated script that segments cells using Cellpose ^85^ and aligns them at each time frame based on their centroids. Photobleaching correction was applied using the bleach-correct Napari plugin. We then selected the spot detection threshold and cluster detection parameters for each cell. Using the spot and Cluster Detection module of bigFishLive, we obtained the nucleoplasmic spots for each frame. The detected spots were used to build a reference spot which models the signal from a single molecule of RNA. This reference spot was then used to decompose the transcription site, into a composite number of single RNA molecules. To track transcription sites and obtain their trajectories, we pre-detected their position in each frame using a computational blurring technique that removed single molecules. The tracks were manually corrected and used to obtain high confidence transcription site signals. Finally, we calculated the exact number of nascent RNA over time at each transcription site.

### Integrating MS2 and SMF data using mathematical modeling

We introduced a novel method to integrate MS2 and SMF data. In this method, transcription dynamics is modeled using *N* state, continuous-time Markov chains with *N*=4, or *N*=5. The model includes the kinetic rate constant vector ***k***, which represent transition rates between states.

Using BurstDeconv ^59^, we deconvolved the single cell MS2 signal and obtained the sequence of transcription non-abortive initiation events from which we computed the empirical survival function. Notably, the advantage of this method is that the calculation of predicted survival functions, occupancy probabilities, and physical constraints does not require time-consuming simulations of the Markov chain model, as our general framework provides analytical formulas for any model. This enables testing a large number of models.

The survival function is defined as

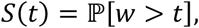

where *w* is the waiting time between successive events, and is estimated using the Kaplan-Meier method.

The deconvolution of the initiation events and the estimate of the survival function are performed using long movies (12 h) with a time resolution of 1 min. Like in ^48,59,60^, the survival function is modeled as a multi-exponential distribution

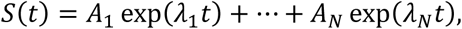

where the coefficients *A_i_* have unit sum,

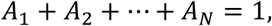

and 𝜆_𝑖_ are negative parameters representing eigenvalues of the Markov chain transition matrix.

The SMF data was used to obtain experimental values of the probabilities *p_1_*, *p_2_*, …, *p_N_* to be in different promoter states. Finally, we also used smFISH to estimate the mean RNA production.

The model parameters 𝒌 are the transition rates between the promoter states and the transcription initiation rate in the productive state. They are the elements of a matrix 𝑸(𝒌) (see Supplemental Methods and ^60^. The characteristic polynomial of 𝑸(𝒌) is

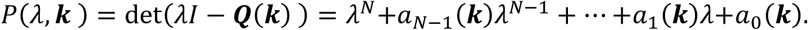

The zeros of 𝑃(𝜆, 𝒌) should be equal to the exponents 𝜆_1_, …, 𝜆_𝑁_.

The model parameters are constrained by the data in two ways. First, the Markov chain model predicts values of the occupancy probabilities as functions of the kinetic parameters (see Supplemental Methods and ^60^)

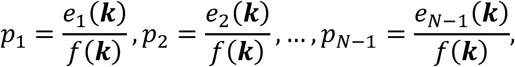

where *e_i_(**k**)* and *f(**k**)* are multivariate polynomials in the kinetic rate constants ***k*** (see Supplemental Methods). These relations can be also used to predict the mean RNA production rate as

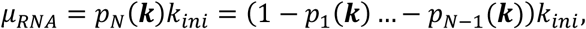

where *k*_ini_ is the transcription initiation rate in the productive state and *p_N_*is the probability of the productive state *N* (considered the unique productive state in these models).

Second, the model also predicts algebraic relations between the parameters 𝑨 = (𝐴_1_, …, 𝐴_𝑁_), 𝝀 = (𝜆_1_, …, 𝜆_𝑁_) of the survival function and the kinetic rate constants ***k***. These constraints are of two types, one type resulting from the Vieta formulas applied to eigenvalues and the second type resulting from eigenvectors of the matrix 𝑸(𝒌).

All the constraints are taken into account as penalty terms in the objective function

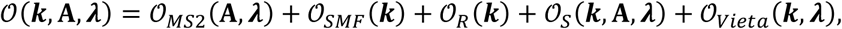

where

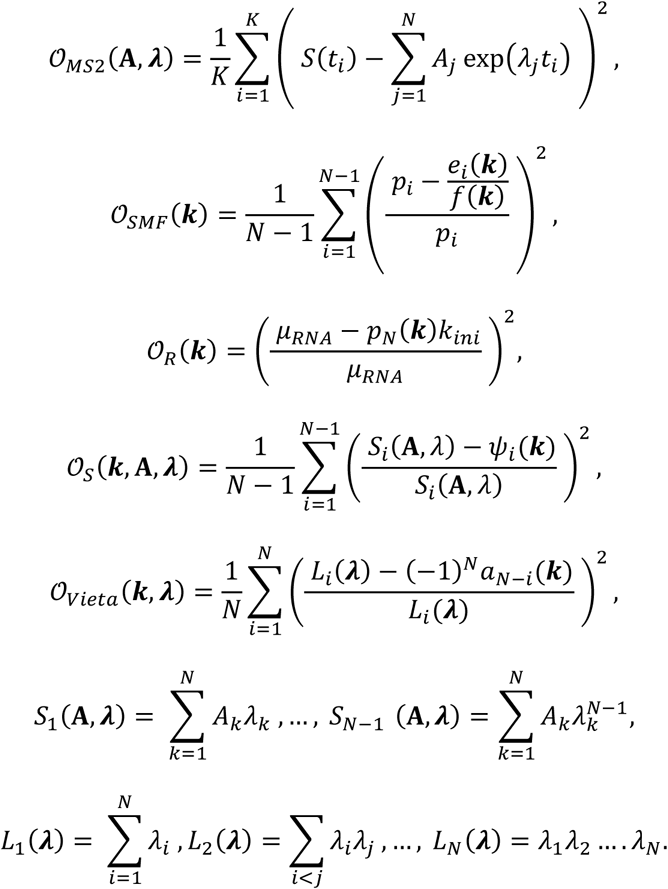

The objective function 𝒪(𝒌, 𝐀, 𝝀) was minimized with respect to the parameters 𝒌, 𝐀, 𝝀. The optimization was performed using gradient descent in Matlab using a multi-start local optimization method with logarithmic sampling of the start values. Parametric uncertainty was assessed by analyzing the parameter values for both optimal and sub-optimal solutions, resulting from various starting parameter values.

## Acknowledgements

We thank Monsef Benkirane for useful discussions and Shinichi Machida for help with the capture-seq MNase experiment. We acknowledge and thank members of the MRI imaging facility, part of the national infrastructure France-BioImaging supported by the French National Research Agency (ANR-10-INBS-04, ‘Investissement d’Avenir’ program). Sequencing experiments were performed by Montpellier GenomiX facility (MGX), which acknowledges financial support from France Génomique National Infrastructure, funded as part of ‘Investissement d’Avenir’ program managed by the French National Research Agency (ANR-10-INBS-09). V.S and F.M. were supported by fellowships from the Agence Nationale de la Recherche sur le SIDA (ANRS-MIE). V.S. was also supported by a fellowship from SIDACTION. The work was supported by grants from the ANRS-MIE and SIDACTION to E.B.

**Supplementary Figure : see attached file**

## References

1. Pichon, X., Lagha, M., Mueller, F., and Bertrand, E. (2018). A Growing Toolbox to Image Gene Expression in Single Cells: Sensitive Approaches for Demanding Challenges. Mol Cell 71, 468–480. 10.1016/j.molcel.2018.07.022.

2. Chubb, J.R., Trcek, T., Shenoy, S.M., and Singer, R.H. (2006). Transcriptional pulsing of a developmental gene. Curr Biol 16, 1018–1025. 10.1016/j.cub.2006.03.092.

3. Whitney, P.H., and Lionnet, T. (2024). The method in the madness: Transcriptional control from stochastic action at the single-molecule scale. Curr Opin Struct Biol 87, 102873. 10.1016/j.sbi.2024.102873.

4. Rodriguez, J., and Larson, D.R. (2020). Transcription in Living Cells: Molecular Mechanisms of Bursting. Annu Rev Biochem 89, 189–212. 10.1146/annurev-biochem-011520-105250.

5. Tunnacliffe, E., and Chubb, J.R. (2020). What Is a Transcriptional Burst? Trends Genet 36, 288–297. 10.1016/j.tig.2020.01.003.

6. Tantale, K., Mueller, F., Kozulic-Pirher, A., Lesne, A., Victor, J.-M., Robert, M.-C., Capozi, S., Chouaib, R., Bäcker, V., Mateos-Langerak, J., et al. (2016). A single-molecule view of transcription reveals convoys of RNA polymerases and multi-scale bursting. Nat Commun 7, 12248. 10.1038/ncomms12248.

7. Corrigan, A.M., Tunnacliffe, E., Cannon, D., and Chubb, J.R. (2016). A continuum model of transcriptional bursting. Elife 5, e13051. 10.7554/eLife.13051.

8. Brown, C.R., Mao, C., Falkovskaia, E., Jurica, M.S., and Boeger, H. (2013). Linking stochastic fluctuations in chromatin structure and gene expression. PLoS Biol 11, e1001621. 10.1371/journal.pbio.1001621.

9. Brown, C.R., and Boeger, H. (2014). Nucleosomal promoter variation generates gene expression noise. Proc Natl Acad Sci U S A 111, 17893–17898. 10.1073/pnas.1417527111.

10. Krebs, A.R., Imanci, D., Hoerner, L., Gaidatzis, D., Burger, L., and Schübeler, D. (2017). Genome-wide Single-Molecule Footprinting Reveals High RNA Polymerase II Turnover at Paused Promoters. Mol Cell 67, 411–422.e4. 10.1016/j.molcel.2017.06.027.

11. Lenstra, T.L., Rodriguez, J., Chen, H., and Larson, D.R. (2016). Transcription Dynamics in Living Cells. Annu Rev Biophys 45, 25–47. 10.1146/annurev-biophys-062215-010838.

12. Meeussen, J.V.W., and Lenstra, T.L. (2024). Time will tell: comparing timescales to gain insight into transcriptional bursting. Trends Genet 40, 160–174. 10.1016/j.tig.2023.11.003.

13. Ahmad, K., Brahma, S., and Henikoff, S. (2024). Epigenetic pioneering by SWI/SNF family remodelers. Mol Cell 84, 194–201. 10.1016/j.molcel.2023.10.045.

14. Kumar, A., Chan, J., Taguchi, M., and Kono, H. (2021). Interplay among transacting factors around promoter in the initial phases of transcription. Curr Opin Struct Biol 71, 7–15. 10.1016/j.sbi.2021.04.008.

15. Gómora-García, J.C., and Furlan-Magaril, M. (2025). Pioneer factors outline chromatin architecture. Curr Opin Cell Biol 93, 102480. 10.1016/j.ceb.2025.102480.

16. Kornberg, R.D., and Lorch, Y. (2020). Primary Role of the Nucleosome. Mol Cell 79, 371–375. 10.1016/j.molcel.2020.07.020.

17. Malik, S., and Roeder, R.G. (2023). Regulation of the RNA polymerase II pre-initiation complex by its associated coactivators. Nat Rev Genet 24, 767–782. 10.1038/s41576-023-00630-9.

18. Sun, R., and Fisher, R.P. (2025). The CDK9-SPT5 Axis in Control of Transcription Elongation by RNAPII. J Mol Biol 437, 168746. 10.1016/j.jmb.2024.168746.

19. Chang, H.H., Hemberg, M., Barahona, M., Ingber, D.E., and Huang, S. (2008). Transcriptome-wide noise controls lineage choice in mammalian progenitor cells. Nature 453, 544–547. 10.1038/nature06965.

20. Raj, A., Rifkin, S.A., Andersen, E., and van Oudenaarden, A. (2010). Variability in gene expression underlies incomplete penetrance. Nature 463, 913–918. 10.1038/nature08781.

21. Wernet, M.F., Mazzoni, E.O., Çelik, A., Duncan, D.M., Duncan, I., and Desplan, C. (2006). Stochastic spineless expression creates the retinal mosaic for colour vision. Nature 440, 174–180. 10.1038/nature04615.

22. Ho, Y.-C., Shan, L., Hosmane, N.N., Wang, J., Laskey, S.B., Rosenbloom, D.I.S., Lai, J., Blankson, J.N., Siliciano, J.D., and Siliciano, R.F. (2013). Replication-competent noninduced proviruses in the latent reservoir increase barrier to HIV-1 cure. Cell 155, 540–551. 10.1016/j.cell.2013.09.020.

23. Weinberger, L.S., Burnett, J.C., Toettcher, J.E., Arkin, A.P., and Schaffer, D.V. (2005). Stochastic gene expression in a lentiviral positive-feedback loop: HIV-1 Tat fluctuations drive phenotypic diversity. Cell 122, 169–182. 10.1016/j.cell.2005.06.006.

24. Rouzine, I.M., Razooky, B.S., and Weinberger, L.S. (2014). Stochastic variability in HIV affects viral eradication. Proc Natl Acad Sci U S A 111, 13251–13252. 10.1073/pnas.1413362111.

25. Damour, A., Slaninova, V., Radulescu, O., Bertrand, E., and Basyuk, E. (2023). Transcriptional Stochasticity as a Key Aspect of HIV-1 Latency. Viruses 15, 1969. 10.3390/v15091969.

26. Dufour, C., Gantner, P., Fromentin, R., and Chomont, N. (2020). The multifaceted nature of HIV latency. J Clin Invest 130, 3381–3390. 10.1172/JCI136227.

27. Cary, D.C., Fujinaga, K., and Peterlin, B.M. (2016). Molecular mechanisms of HIV latency. J Clin Invest 126, 448–454. 10.1172/JCI80565.

28. Laird, G.M., Bullen, C.K., Rosenbloom, D.I.S., Martin, A.R., Hill, A.L., Durand, C.M., Siliciano, J.D., and Siliciano, R.F. (2015). Ex vivo analysis identifies effective HIV-1 latency-reversing drug combinations. J Clin Invest 125, 1901–1912. 10.1172/JCI80142.

29. Berkhout, B., and Jeang, K.T. (1992). Functional roles for the TATA promoter and enhancers in basal and Tat-induced expression of the human immunodeficiency virus type 1 long terminal repeat. J Virol 66, 139–149. 10.1128/JVI.66.1.139-149.1992.

30. Kao, S.Y., Calman, A.F., Luciw, P.A., and Peterlin, B.M. (1987). Anti-termination of transcription within the long terminal repeat of HIV-1 by tat gene product. Nature 330, 489–493. 10.1038/330489a0.

31. Ping, Y.H., and Rana, T.M. (2001). DSIF and NELF interact with RNA polymerase II elongation complex and HIV-1 Tat stimulates P-TEFb-mediated phosphorylation of RNA polymerase II and DSIF during transcription elongation. J Biol Chem 276, 12951–12958. 10.1074/jbc.M006130200.

32. Su, B.G., and Vos, S.M. (2024). Distinct negative elongation factor conformations regulate RNA polymerase II promoter-proximal pausing. Mol Cell 84, 1243–1256.e5. 10.1016/j.molcel.2024.01.023.

33. Vos, S.M., Farnung, L., Urlaub, H., and Cramer, P. (2018). Structure of paused transcription complex Pol II-DSIF-NELF. Nature 560, 601–606. 10.1038/s41586-018-0442-2.

34. Wei, P., Garber, M.E., Fang, S.M., Fischer, W.H., and Jones, K.A. (1998). A novel CDK9-associated C-type cyclin interacts directly with HIV-1 Tat and mediates its high-affinity, loop-specific binding to TAR RNA. Cell 92, 451–462. 10.1016/s0092-8674(00)80939-3.

35. Sobhian, B., Laguette, N., Yatim, A., Nakamura, M., Levy, Y., Kiernan, R., and Benkirane, M. (2010). HIV-1 Tat assembles a multifunctional transcription elongation complex and stably associates with the 7SK snRNP. Mol Cell 38, 439–451. 10.1016/j.molcel.2010.04.012.

36. Schulze-Gahmen, U., and Hurley, J.H. (2018). Structural mechanism for HIV-1 TAR loop recognition by Tat and the super elongation complex. Proc Natl Acad Sci U S A 115, 12973–12978. 10.1073/pnas.1806438115.

37. He, N., Liu, M., Hsu, J., Xue, Y., Chou, S., Burlingame, A., Krogan, N.J., Alber, T., and Zhou, Q. (2010). HIV-1 Tat and host AFF4 recruit two transcription elongation factors into a bifunctional complex for coordinated activation of HIV-1 transcription. Mol Cell 38, 428–438. 10.1016/j.molcel.2010.04.013.

38. Vos, S.M., Farnung, L., Boehning, M., Wigge, C., Linden, A., Urlaub, H., and Cramer, P. (2018). Structure of activated transcription complex Pol II-DSIF-PAF-SPT6. Nature 560, 607–612. 10.1038/s41586-018-0440-4.

39. Stadelmayer, B., Micas, G., Gamot, A., Martin, P., Malirat, N., Koval, S., Raffel, R., Sobhian, B., Severac, D., Rialle, S., et al. (2014). Integrator complex regulates NELF-mediated RNA polymerase II pause/release and processivity at coding genes. Nat Commun 5, 5531. 10.1038/ncomms6531.

40. Fianu, I., Ochmann, M., Walshe, J.L., Dybkov, O., Cruz, J.N., Urlaub, H., and Cramer, P. (2024). Structural basis of Integrator-dependent RNA polymerase II termination. Nature 629, 219–227. 10.1038/s41586-024-07269-4.

41. Marzio, G., Tyagi, M., Gutierrez, M.I., and Giacca, M. (1998). HIV-1 tat transactivator recruits p300 and CREB-binding protein histone acetyltransferases to the viral promoter. Proc Natl Acad Sci U S A 95, 13519–13524. 10.1073/pnas.95.23.13519.

42. Benkirane, M., Chun, R.F., Xiao, H., Ogryzko, V.V., Howard, B.H., Nakatani, Y., and Jeang, K.T. (1998). Activation of integrated provirus requires histone acetyltransferase. p300 and P/CAF are coactivators for HIV-1 Tat. J Biol Chem 273, 24898–24905. 10.1074/jbc.273.38.24898.

43. De Crignis, E., and Mahmoudi, T. (2017). The Multifaceted Contributions of Chromatin to HIV-1 Integration, Transcription, and Latency. Int Rev Cell Mol Biol 328, 197–252. 10.1016/bs.ircmb.2016.08.006.

44. Mizutani, T., Ishizaka, A., Tomizawa, M., Okazaki, T., Yamamichi, N., Kawana-Tachikawa, A., Iwamoto, A., and Iba, H. (2009). Loss of the Brm-type SWI/SNF chromatin remodeling complex is a strong barrier to the Tat-independent transcriptional elongation of human immunodeficiency virus type 1 transcripts. J Virol 83, 11569–11580. 10.1128/JVI.00742-09.

45. Jones, J.E., Gunderson, C.E., Wigdahl, B., and Nonnemacher, M.R. (2025). Impact of chromatin on HIV-1 latency: a multi-dimensional perspective. Epigenetics Chromatin 18, 9. 10.1186/s13072-025-00573-x.

46. Lusic, M., Marcello, A., Cereseto, A., and Giacca, M. (2003). Regulation of HIV-1 gene expression by histone acetylation and factor recruitment at the LTR promoter. EMBO J 22, 6550–6561. 10.1093/emboj/cdg631.

47. Rafati, H., Parra, M., Hakre, S., Moshkin, Y., Verdin, E., and Mahmoudi, T. (2011). Repressive LTR nucleosome positioning by the BAF complex is required for HIV latency. PLoS Biol 9, e1001206. 10.1371/journal.pbio.1001206.

48. Tantale, K., Garcia-Oliver, E., Robert, M.-C., L’Hostis, A., Yang, Y., Tsanov, N., Topno, R., Gostan, T., Kozulic-Pirher, A., Basu-Shrivastava, M., et al. (2021). Stochastic pausing at latent HIV-1 promoters generates transcriptional bursting. Nat Commun 12, 4503. 10.1038/s41467-021-24462-5.

49. Kleinendorst, R.W.D., Barzaghi, G., Smith, M.L., Zaugg, J.B., and Krebs, A.R. (2021). Genome-wide quantification of transcription factor binding at single-DNA-molecule resolution using methyl-transferase footprinting. Nat Protoc 16, 5673–5706. 10.1038/s41596-021-00630-1.

50. Bertrand, E., Chartrand, P., Schaefer, M., Shenoy, S.M., Singer, R.H., and Long, R.M. (1998). Localization of ASH1 mRNA particles in living yeast. Mol Cell 2, 437–445. 10.1016/s1097-2765(00)80143-4.

51. Fusco, D., Accornero, N., Lavoie, B., Shenoy, S.M., Blanchard, J.-M., Singer, R.H., and Bertrand, E. (2003). Single mRNA molecules demonstrate probabilistic movement in living mammalian cells. Curr Biol 13, 161–167. 10.1016/s0960-9822(02)01436-7.

52. Dalakishvili, L., Managori, H., Bardet, A., Slaninová, V., Bertrand, E., and Molina, N. (2025). Biophysical Modeling Uncovers Transcription Factor and Nucleosome Binding on Single DNA Molecules. Preprint at Genomics, 10.1101/2025.05.13.653852 https://doi.org/10.1101/2025.05.13.653852.

53. Rauluseviciute, I., Riudavets-Puig, R., Blanc-Mathieu, R., Castro-Mondragon, J.A., Ferenc, K., Kumar, V., Lemma, R.B., Lucas, J., Chèneby, J., Baranasic, D., et al. (2024). JASPAR 2024: 20th anniversary of the open-access database of transcription factor binding profiles. Nucleic Acids Res 52, D174–D182. 10.1093/nar/gkad1059.

54. Pachkov, M., Erb, I., Molina, N., and van Nimwegen, E. (2007). SwissRegulon: a database of genome-wide annotations of regulatory sites. Nucleic Acids Res 35, D127–131. 10.1093/nar/gkl857.

55. Verdin, E., Paras, P., and Van Lint, C. (1993). Chromatin disruption in the promoter of human immunodeficiency virus type 1 during transcriptional activation. EMBO J 12, 3249–3259. 10.1002/j.1460-2075.1993.tb05994.x.

56. Raha, T., Cheng, S.W.G., and Green, M.R. (2005). HIV-1 Tat stimulates transcription complex assembly through recruitment of TBP in the absence of TAFs. PLoS Biol 3, e44. 10.1371/journal.pbio.0030044.

57. Femino, A.M., Fay, F.S., Fogarty, K., and Singer, R.H. (1998). Visualization of single RNA transcripts in situ. Science 280, 585–590. 10.1126/science.280.5363.585.

58. Vansant, G., Bruggemans, A., Janssens, J., and Debyser, Z. (2020). Block-And-Lock Strategies to Cure HIV Infection. Viruses 12, 84. 10.3390/v12010084.

59. Douaihy, M., Topno, R., Lagha, M., Bertrand, E., and Radulescu, O. (2023). BurstDECONV: a signal deconvolution method to uncover mechanisms of transcriptional bursting in live cells. Nucleic Acids Res 51, e88. 10.1093/nar/gkad629.

60. Radulescu, O., Grigoriev, D., Seiss, M., Douaihy, M., Lagha, M., and Bertrand, E. (2024). Identifying Markov Chain Models from Time-to-Event Data: An Algebraic Approach. Bull Math Biol 87, 11. 10.1007/s11538-024-01385-y.

61. Machida, S., Depierre, D., Chen, H.-C., Thenin-Houssier, S., Petitjean, G., Doyen, C.M., Takaku, M., Cuvier, O., and Benkirane, M. (2020). Exploring histone loading on HIV DNA reveals a dynamic nucleosome positioning between unintegrated and integrated viral genome. Proc Natl Acad Sci U S A 117, 6822–6830. 10.1073/pnas.1913754117.

62. Geis, F.K., and Goff, S.P. (2019). Unintegrated HIV-1 DNAs are loaded with core and linker histones and transcriptionally silenced. Proc Natl Acad Sci U S A 116, 23735– 23742. 10.1073/pnas.1912638116.

63. Schlesinger, S., Kaffe, B., Melcer, S., Aguilera, J.D., Sivaraman, D.M., Kaplan, T., and Meshorer, E. (2017). A hyperdynamic H3.3 nucleosome marks promoter regions in pluripotent embryonic stem cells. Nucleic Acids Res 45, 12181–12194. 10.1093/nar/gkx817.

64. Dion, M.F., Kaplan, T., Kim, M., Buratowski, S., Friedman, N., and Rando, O.J. (2007). Dynamics of replication-independent histone turnover in budding yeast. Science 315, 1405–1408. 10.1126/science.1134053.

65. Kraushaar, D.C., Jin, W., Maunakea, A., Abraham, B., Ha, M., and Zhao, K. (2013). Genome-wide incorporation dynamics reveal distinct categories of turnover for the histone variant H3.3. Genome Biol 14, R121. 10.1186/gb-2013-14-10-r121.

66. Deal, R.B., Henikoff, J.G., and Henikoff, S. (2010). Genome-wide kinetics of nucleosome turnover determined by metabolic labeling of histones. Science 328, 1161– 1164. 10.1126/science.1186777.

67. De Graaf, P., Mousson, F., Geverts, B., Scheer, E., Tora, L., Houtsmuller, A.B., and Timmers, H.Th.M. (2010). Chromatin interaction of TATA-binding protein is dynamically regulated in human cells. Journal of Cell Science 123, 2663–2671. 10.1242/jcs.064097.

68. Szczurek, A.T., Dimitrova, E., Kelley, J.R., Blackledge, N.P., and Klose, R.J. (2024). The Polycomb system sustains promoters in a deep OFF state by limiting pre-initiation complex formation to counteract transcription. Nat Cell Biol 26, 1700–1711. 10.1038/s41556-024-01493-w.

69. Hasegawa, Y., and Struhl, K. (2019). Promoter-specific dynamics of TATA-binding protein association with the human genome. Genome Res 29, 1939–1950. 10.1101/gr.254466.119.

70. Nguyen, V.Q., Ranjan, A., Liu, S., Tang, X., Ling, Y.H., Wisniewski, J., Mizuguchi, G., Li, K.Y., Jou, V., Zheng, Q., et al. (2021). Spatiotemporal coordination of transcription preinitiation complex assembly in live cells. Mol Cell 81, 3560–3575.e6. 10.1016/j.molcel.2021.07.022.

71. Forero-Quintero, L.S., Raymond, W., Handa, T., Saxton, M.N., Morisaki, T., Kimura, H., Bertrand, E., Munsky, B., and Stasevich, T.J. (2021). Live-cell imaging reveals the spatiotemporal organization of endogenous RNA polymerase II phosphorylation at a single gene. Nat Commun 12, 3158. 10.1038/s41467-021-23417-0.

72. Erickson, B., Sheridan, R.M., Cortazar, M., and Bentley, D.L. (2018). Dynamic turnover of paused Pol II complexes at human promoters. Genes Dev 32, 1215–1225. 10.1101/gad.316810.118.

73. Henriques, T., Gilchrist, D.A., Nechaev, S., Bern, M., Muse, G.W., Burkholder, A., Fargo, D.C., and Adelman, K. (2013). Stable pausing by RNA polymerase II provides an opportunity to target and integrate regulatory signals. Mol Cell 52, 517–528. 10.1016/j.molcel.2013.10.001.

74. Taft, R.J., Glazov, E.A., Cloonan, N., Simons, C., Stephen, S., Faulkner, G.J., Lassmann, T., Forrest, A.R.R., Grimmond, S.M., Schroder, K., et al. (2009). Tiny RNAs associated with transcription start sites in animals. Nat Genet 41, 572–578. 10.1038/ng.312.

75. Shelansky, R., Abrahamsson, S., Doody, M., Brown, C.R., Patel, H.P., Lenstra, T.L., Larson, D.R., and Boeger, H. (2022). A Telltale Sign of Irreversibility in Transcriptional Regulation. Preprint at Molecular Biology, 10.1101/2022.06.27.497819 https://doi.org/10.1101/2022.06.27.497819.

76. Garland, W., and Jensen, T.H. (2024). Nuclear sorting of short RNA polymerase II transcripts. Mol Cell 84, 3644–3655. 10.1016/j.molcel.2024.08.024.

77. Hughes, A.L., Szczurek, A.T., Kelley, J.R., Lastuvkova, A., Turberfield, A.H., Dimitrova, E., Blackledge, N.P., and Klose, R.J. (2023). A CpG island-encoded mechanism protects genes from premature transcription termination. Nat Commun 14, 726. 10.1038/s41467-023-36236-2.

78. Matkovic, R., Morel, M., Lanciano, S., Larrous, P., Martin, B., Bejjani, F., Vauthier, V., Hansen, M.M.K., Emiliani, S., Cristofari, G., et al. (2022). TASOR epigenetic repressor cooperates with a CNOT1 RNA degradation pathway to repress HIV. Nat Commun 13, 66. 10.1038/s41467-021-27650-5.

79. Ajit, K., Alagia, A., Burger, K., and Gullerova, M. (2024). Tyrosine 1-phosphorylated RNA polymerase II transcribes PROMPTs to facilitate proximal promoter pausing and induce global transcriptional repression in response to DNA damage. Genome Res 34, 201–216. 10.1101/gr.278644.123.

80. Lee, J.-H., and Skalnik, D.G. (2008). Wdr82 is a C-terminal domain-binding protein that recruits the Setd1A Histone H3-Lys4 methyltransferase complex to transcription start sites of transcribed human genes. Mol Cell Biol 28, 609–618. 10.1128/MCB.01356-07.

81. Materne, P., Anandhakumar, J., Migeot, V., Soriano, I., Yague-Sanz, C., Hidalgo, E., Mignion, C., Quintales, L., Antequera, F., and Hermand, D. (2015). Promoter nucleosome dynamics regulated by signalling through the CTD code. Elife 4, e09008. 10.7554/eLife.09008.

82. Zuker, M. (2003). Mfold web server for nucleic acid folding and hybridization prediction. Nucleic Acids Res 31, 3406–3415. 10.1093/nar/gkg595.

83. Tsanov, N., Samacoits, A., Chouaib, R., Traboulsi, A.-M., Gostan, T., Weber, C., Zimmer, C., Zibara, K., Walter, T., Peter, M., et al. (2016). smiFISH and FISH-quant-a flexible single RNA detection approach with super-resolution capability. Nucleic Acids Res 44, e165. 10.1093/nar/gkw784.

84. Imbert, A., Ouyang, W., Safieddine, A., Coleno, E., Zimmer, C., Bertrand, E., Walter, T., and Mueller, F. (2022). FISH-quant v2: a scalable and modular tool for smFISH image analysis. RNA 28, 786–795. 10.1261/rna.079073.121.

85. Stringer, C., Wang, T., Michaelos, M., and Pachitariu, M. (2021). Cellpose: a generalist algorithm for cellular segmentation. Nat Methods 18, 100–106. 10.1038/s41592-020-01018-x.

